# Astrocyte-Guided Maturation of Neural Constructs in a Modular Biosynthetic Hydrogel for Biohybrid Neurotechnologies

**DOI:** 10.1101/2025.08.22.671553

**Authors:** Martina Genta, Sofia Peressotti, Roberto Portillo-Lara, Josef Goding, Rylie Green

**Affiliations:** Bioengineering Department, Imperial College London, South Kensington, London, SW7 2AZ, UK; Psychiatry and Fundamental Neuroscience Department, University of Geneva, 1202, Switzerland

## Abstract

Bionic implants are increasingly used to restore neural function yet achieving a chronically stable neural interface remains challenging. Biohybrid neurotechnologies aim to overcome this limitation by integrating living tissue components that can promote long-term performance and functional integration with the nervous system. However, existing biomaterial coatings often lack the physiological complexity, cellular heterogeneity, and neurotrophic support required to sustain neural network formation. Here, we present a modular strategy which leverages astrocyte-guided mechanisms of neural network development. A biosynthetic hydrogel was developed using norbornene-functionalized poly(vinyl alcohol) and gelatin (PVA-GEL), crosslinked via visible light-triggered thiol–ene chemistry, to yield bioactive and highly tuneable scaffolds. A systematic characterization varying gelatin content and polymer weight enabled the identification an optimal formulation to promote astrocytic growth, while maintaining mechanical stability and degradation profiles that are compatible with brain implants. Co-encapsulation with neural progenitors promoted neuronal differentiation, neurite outgrowth, and the formation of synaptically competent networks. The construct developed into functional interfaces when in contact with brain tissue *ex vivo*, highlighting its potential as a biohybrid electrode coating. This work lays the foundation for the development of biologically guided biohybrid interfaces, towards seamless integration with the nervous system.

## 1. Introduction

Neural implants are widely used in the management of a variety of neurological and non-neurological conditions, as well as the restoration of sensorimotor and autonomic functions impaired by trauma or disease^1,2^. However, their performance is often hindered by an inflammatory foreign body reaction triggered upon implantation, which ultimately leads to fibrotic scar formation and poor device performance across chronic time frames^3–5^. To address this limitation, multiple biomimetic design strategies have been explored based on recapitulating the physical properties, form factor, composition, and microarchitecture of neural tissues^6–8^. More recently, the exploration of tissue engineering approaches has led to the design of biohybrid neural interfaces which incorporate living cells in or around the device, to promote biointegration and enable biologically mediated communication with the nervous system^9–12^. This approach could enable direct synaptic interfacing with the host nervous system, enabling long-term performance and more selective neuromodulation, while also reducing inflammatory responses triggered by conventional implant designs^3,4,13^. However, engineering structurally permissive and bioinstructive materials that support the formation, maturation, and functional integration of tissue-engineered constructs for neural applications remains a significant challenge.

Hydrogels are well suited for this application due to their tuneable physical properties which can emulate the native extracellular matrix (ECM) and their capacity to deliver bio-instructive cues to guide the development of neural constructs^14–17^. In the context of neural interfaces, three main aspects should be considered when designing biomaterial systems for the fabrication of cell and tissue culture scaffolds. First, the physical properties and degradation profiles of hydrogel coatings should be finely tuned for cell support and device integration. Although complete biodegradation is optimal within the context of implantable biomaterials, the rate at which this process occurs should allow encapsulated cells to develop and remodel the material construct during the first weeks post-implantation^18^. Thus, material composition and programmable, cell-driven biodegradability are key parameters of scaffold design that greatly influence cell growth and spatial organization, while maintaining the stability of the tissue-engineered interface. Second, metallic electrodes are remarkably stiff compared to neural tissue, which enhances inflammation and foreign body response^19^. Other mechanical properties, such as viscoelasticity, are also known to greatly influence cell responses and adhesion^20^. Thus, material-based approaches should recapitulate the mechanical properties of brain tissue. Third, reproducing the intricate crosstalk between neurons and glial cells is essential to ensure adequate construct maturation and functionality. Astrocytes are anchorage-dependent and require a supportive 3D architecture that mimics the mechanical properties of the ECM^21,22^ to survive, grow, and develop^23–25^. They play key roles in synaptogenesis, plasticity, neuronal signaling and neural progenitor cell (NPC) fate via trophic support and ECM remodelling^26–29^. The incorporation of cell adhesive motifs, such as the arginine-glycine-aspartic acid (RGD) epitope, is a powerful strategy to achieve functional cell-material interactions. ECM remodelling is achieved by secretion of matrix metalloproteinase (MMP) enzymes, which degrade MMP-sensitive molecules in their natural or synthetic environment^30^. As the physical and biological properties of scaffolds have a profound influence on astrocytic phenotype^31,32^, designing hydrogel scaffolds that effectively support astrocyte growth and function could constitute an indirect but powerful strategy to engineer robust neural constructs for a variety of applications.

Co-culture systems comprising neurons and astrocytes are extensively used to replicate the multicellular organisation of the CNS for fundamental and translational research^33–37^. 3D-encapsulation into hydrogel scaffolds has also been widely explored to generate physiologically relevant environments for studies of neural development, function, and disease^38,39^. Although several studies have focused on embedding combinations of terminally differentiated neuronal and glial phenotypes into soft matrices^38^, this approach bypasses the endogenous processes that underlie tissue development and morphogenesis. Alternatively, leveraging the natural interplay between NPCs and astrocytes could better recapitulate the endogenous cues that drive neural differentiation and synaptic maturation. One salient example of this approach is the living electrode platform developed by Goding et al.^11^, which consists in microelectrode arrays with multi-layered tissue-engineered coatings. The layered construct is composed of a PVA-functionalized hydrogel embedded with neural cells, which is formed on top of an electroconductive PVA-based hydrogel in direct contact with the electrode substrate. The conductive hydrogel layer is known to improve the mechanical mismatch with the tissue and the electrical performance of the device, while protecting the encapsulated cells from the high-voltage capacitive charging that may occur on metallic electrodes^11,12,40^. It is hypothesised that this biohybrid approach could ameliorate the foreign body reaction and promote functional integration of neural interfaces into the host nervous system upon implantation. However, despite having no negative effects on electrode performance, the limited cell development within the biosynthetic layer (PVA functionalized with 10% sericin and gelatin) compromised the viability, neural cell growth and differentiation within the construct in a recent study^41^. This was attributed to the low biological functionalization, which was initially designed to maximize the mechanical stability of the material, but it did not provide the necessary support to the glial population^41^. In this context, the development of a PVA-based hydrogel with tailorable properties and high biological content is essential to advance the living electrode concept, addressing the trade-off between physical stability of the hydrogel layer and bioactive support to neural cells.

Here, a biosynthetic hydrogel system for neural interface coatings is reported to enable the generation of biointegrative and synaptically active neural constructs, by leveraging endogenous glial-mediated mechanisms (**Figure 1**). This biomaterial is based on poly(vinyl alcohol) (PVA) and gelatin (GEL) functionalized with norbornene (NB) groups to enable click-crosslinking via visible light-triggered thiol–ene chemistry. This bio-orthogonal process allows for precise control on polymer composition and minimal off-target interactions with the endogenous microenvironment^30,42^. Boulingre et al.^43^ have recently confirmed the cytocompatibility of the PVA-GEL hydrogel using primary astrocytes and successfully validated its use as a bioactive layer of the living electrode construct. Here, the PVA-GEL hydrogel biophysical properties were thoroughly characterized in multiple polymer compositions (PVA:GEL ratio) and weight percentages, to identify the optimal formulation for brain interface applications and glial cell support. Physical characteristics such as mass swelling ratio, mass loss, degradation profiles and bulk mechanical properties were systematically measured across 28 days of incubation, for a range of polymer compositions. The growth and development of primary astrocytes was then systematically assessed, aiming to generate glial-enriched microenvironments conducive to neuronal development^26,44,45^. This comprehensive biological and physical characterization allowed us to identify an optimal range of physical properties for supporting astrocytes in this material and other applications. Primary astrocytes and NPC-enriched neurospheres were then co-encapsulated within the optimized scaffolds and matured *in vitro* for up to 14 days to generate functional neural constructs. An increased metabolic activity, culture heterogeneity and synaptic expression was observed in the astrocytes-supported cultures compared to neurosphere-only cultures. The interface properties of the PVA-GEL were then investigated by culturing the hydrogels in contact with hippocampal slices *ex vivo*. Tissue ingrowth, cell-cell contacts and synaptic expression was observed in the cultures for chronic time frames (up to 21 days). In summary, PVA-GEL hydrogels could not only be optimized around the complex mechanobiology of glial cells, but also employed to fabricate heterocellular, functional neural constructs that harness the paramount role of astrocytes in neural network development and support.

**Figure 1:**
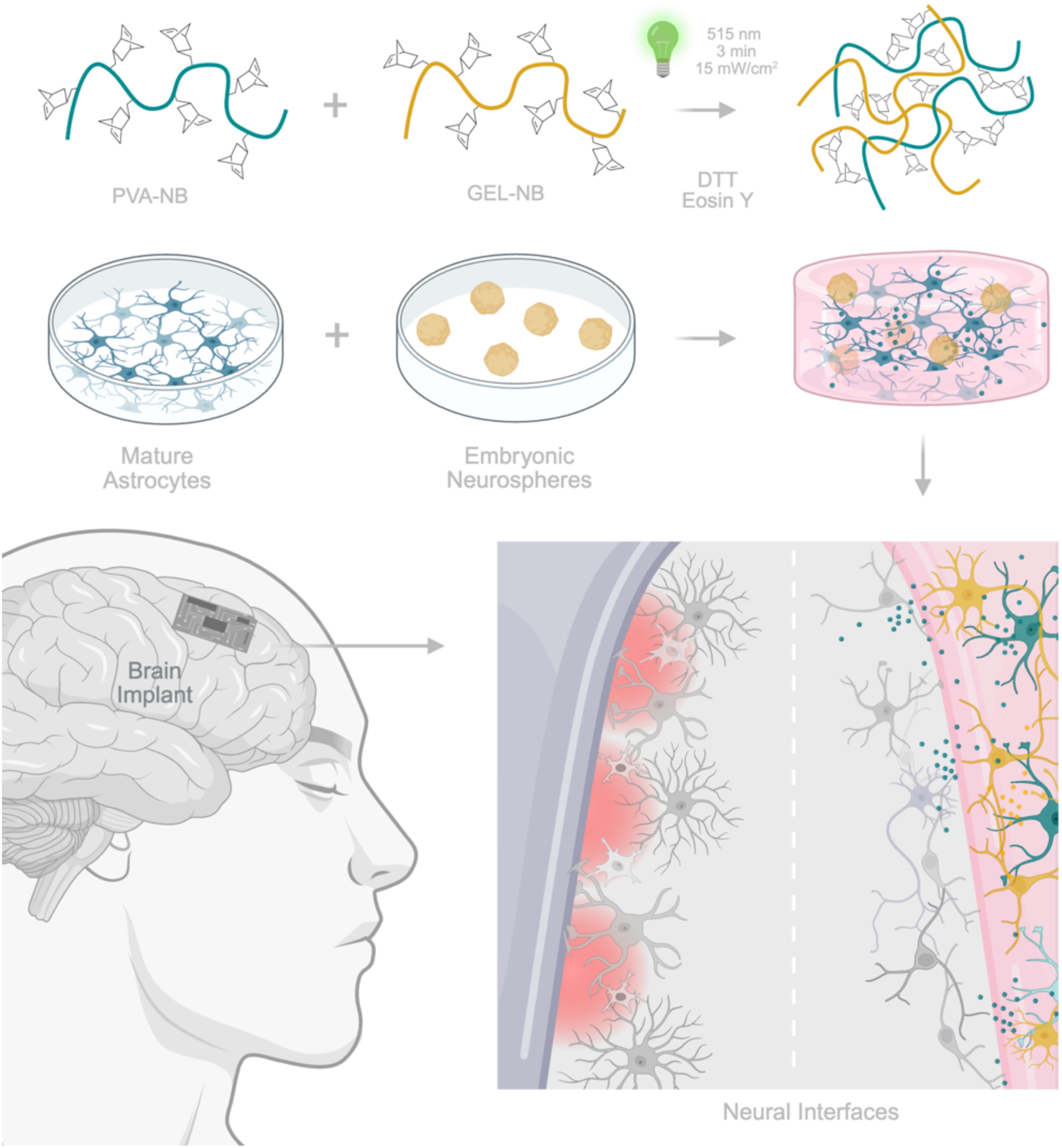
Graphical abstract. A biosynthetic hydrogel material was developed to enhance integration and minimise inflammation in neural interfaces, towards advancing the ‘living electrode concept’. PVA and GEL are first functionalized with norbornene (NB) groups separately and then crosslinked to form the PVA-GEL hydrogel. The PVA functionalization and crosslinking of the hydrogel was achieved via click-chemistry, using dithiothreitol (DTT) as initiator, eosin Y and visible light exposure (515 nm, 3 min, 15 mW/cm^2^). Mature astrocytes and neuronal cultures are co-encapsulated in the PVA-GEL hydrogel. Figure created with Biorender.

This tissue engineering approach holds great potential to fabricate robust biohybrid constructs for the development of living electrodes with enhanced long-term performance and biointegration. Furthermore, this biomimetic approach at generating constructs with tissue-level function and architecture holds great promise for applications in neuroengineering, regenerative medicine, drug screening, and disease modelling.

## 2. Results and Discussion

### 2.1 Polymer Synthesis and Functionalisation

A biosynthetic hydrogel system was engineered based on NB-functionalized PVA and GEL, and its material parameters were systematically optimized to support the growth and development of astrocytes *in vitro*. The synthetic polymer PVA was used as the structural base due to its tuneable mechanical properties, hydrophilicity, and prior use in the fabrication of coatings for neural interfaces^11,46^. The natural polymer GEL was used to impart cell-adhesive and enzymatically degradable properties based on its intrinsic RGD and matrix metalloproteinase (MMP)-sensitive motifs, which are essential to support cell development in tissue-engineered scaffolds^16^. Previous studies have also developed cell-responsive hydrogels by functionalizing GEL with NB groups via click-crosslinking^30,47,48^.

To enable efficient, cytocompatible photo-crosslinking and scaffold modularisation, both PVA and GEL were functionalized with NB groups to allow bio-orthogonal thiol–ene coupling under visible light (**Figure 1**). This crosslinking strategy provides high spatiotemporal control and avoids radical-induced cytotoxicity associated with UV-based polymerisation^49^. A degree of substitution (DS) of 7% was targeted for PVA based on previous literature^50^, and a DS of 18% for GEL to account for its larger molecular weight and to avoid premature gelation from intramolecular crosslinking. Functionalisation of PVA with NB achieved a DS between 5% and 11%, with an overall reaction efficiency of 89.6 ± 3.0% as confirmed by proton nuclear magnetic resonance (¹H-NMR) analysis (**Figure 2A**). GEL-NB was successfully synthesized as confirmed by ^1^H NMR spectra (**Figure 2B**) with an actual DS of 18.9% (**Figure S1**). Photopolymerisation was carried out using DTT as a crosslinker and eosin Y as a photoinitiator, which is activated by visible green light at 515 nm^51^ (**Figure 1**). This crosslinking strategy was used to minimise photodamage and improve compatibility with cells and proteins compared to conventional UV initiation^47,51^. A range of eosin Y concentrations (0.01–1 mM) was tested to optimize the structural stability of the scaffolds without compromising light penetration or generating residual chromophores (**Figure 2C**). Qualitative assessment of the resulting PVA-GEL hydrogels showed that a concentration of 0.1 mM yielded structurally stable and homogeneously crosslinked scaffolds at day 0 and 1 (**Figure 2D**). As potential improvements of the system, Shih et al.^52^ demonstrated that it is possible to use eosin Y alone, improving the ease and cytocompatibility of the crosslinking process.

**Figure 2.**
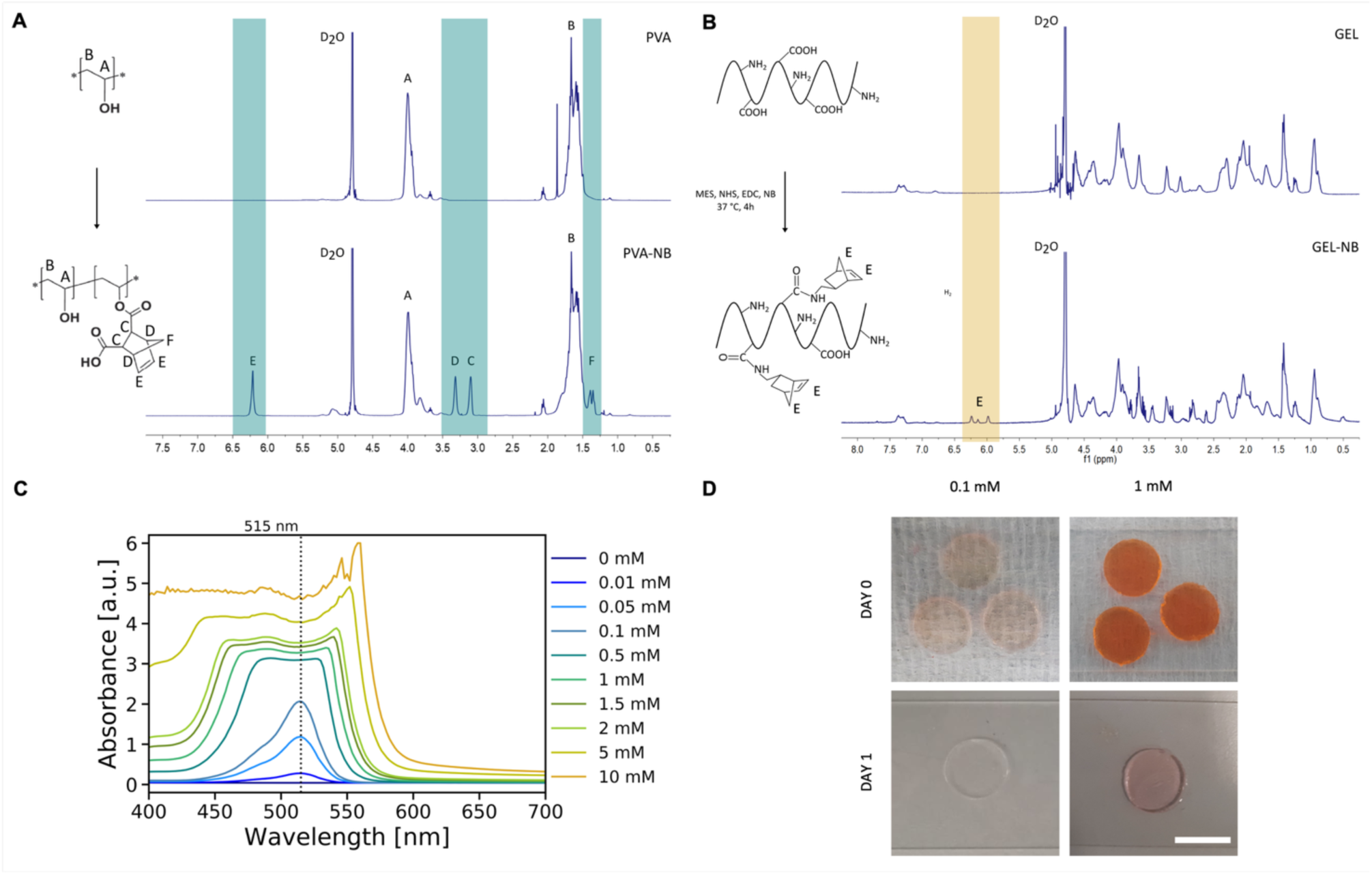
Synthesis and functionalization of the PVA-GEL hydrogel. (A) Chemical characterization of PVA-NB with ^1^H NMR spectra corresponding to pure PVA and conjugated norbornene groups to the PVA backbone (DS data are reported in the supplementary **Figure S1**). (B) Chemical characterization of PVA-NB with ^1^H NMR spectra corresponding to pure GEL and conjugated norbornene groups to the GEL chain. (C) Absorbance spectra of eosin Y at different concentrations in DPBS. (D) Representative images of hydrogels with different concentrations of eosin Y immediately after photopolymerisation (day 0) and after 24 hours of incubation in DPBS (day 1). Scale bar = 10 mm.

The thiol-ene chemistry used here is bio-orthogonal, proceeds rapidly under mild conditions, and provides high specificity, which minimizes the formation of undesired side products^53^. These features are particularly advantageous for encapsulating sensitive cell populations such as astrocytes and NPCs, as they reduce the risk of cytotoxicity and oxidative stress associated with free-radical polymerisation^16^. The modular nature of this strategy also enables fine control over network architecture by tuning the stoichiometry of thiol and ene groups^49^. Furthermore, unreacted NB groups can be leveraged for secondary functionalization with tethering of neurotrophic cues, ECM-mimetic peptides, or bioactive proteins. This in turn enables a variety of multi-modal scaffold customisation strategies, which could be explored to promote specific cellular responses for a broad range of applications^53^.

Taken together, these results demonstrated the successful synthesis of a biosynthetic hydrogel system based on NB-functionalized PVA and GEL, using a tuneable crosslinking strategy based on eosin Y photoinitiation. The physical and mechanical properties of PVA-GEL hydrogels, being key determinants of cell fate and essential for the design of bioinstructive scaffolds, were characterized.

### 2.2 Physical and mechanical characterization

Precise control over the biomechanical and structural properties of hydrogel scaffolds is critical to support cell growth and development in tissue-engineered constructs^54^. Several studies have demonstrated that anchorage-dependent cells such as astrocytes undergo growth arrest and apoptosis when grown without a supportive substrate^55,56^. Cells interact closely with the surrounding ECM, modifying it through complex and dynamic mechanisms. In addition, natural tissues are characterized by specific values of stiffnesses and viscoelastic properties that play a key role in cell development, differentiation and homeostasis^20,57^. Overall, the degradation and mass loss profiles of cell-responsive systems, as well as the material stiffness, matrix viscoelasticity and stress-relaxation properties play an essential role in regulating cell spreading and motility^58^.

The physical properties of PVA-GEL hydrogels were systematically characterized towards identifying scaffold formulations that provided both biological permissiveness and structural robustness compatible with neural interface applications (**Figure 3A**). For this, twelve different hydrogel formulations were produced by varying the ratio of PVA to GEL (100:0, 75:25, 50:50, 25:75) at three polymer weight percentages (5 wt%, 7.5 wt%, and 10 wt%). Hydrogels with 5 wt% were excluded from subsequent mechanical characterization as they degraded after 3-4 days in incubation with DPBS. Pure GEL hydrogels (0:100, PVA:GEL) exhibited poor structural integrity and they were also excluded from further testing. In contrast, mixed formulations produced stable hydrogel networks across a broad range of formulations. FTIR spectroscopy demonstrated successful incorporation of GEL into the PVA network, with increasing gelatin content correlating with stronger amide I absorption at ∼1650 cm⁻¹ ^59,60^ (**Figure 3B**).

**Figure 3.**
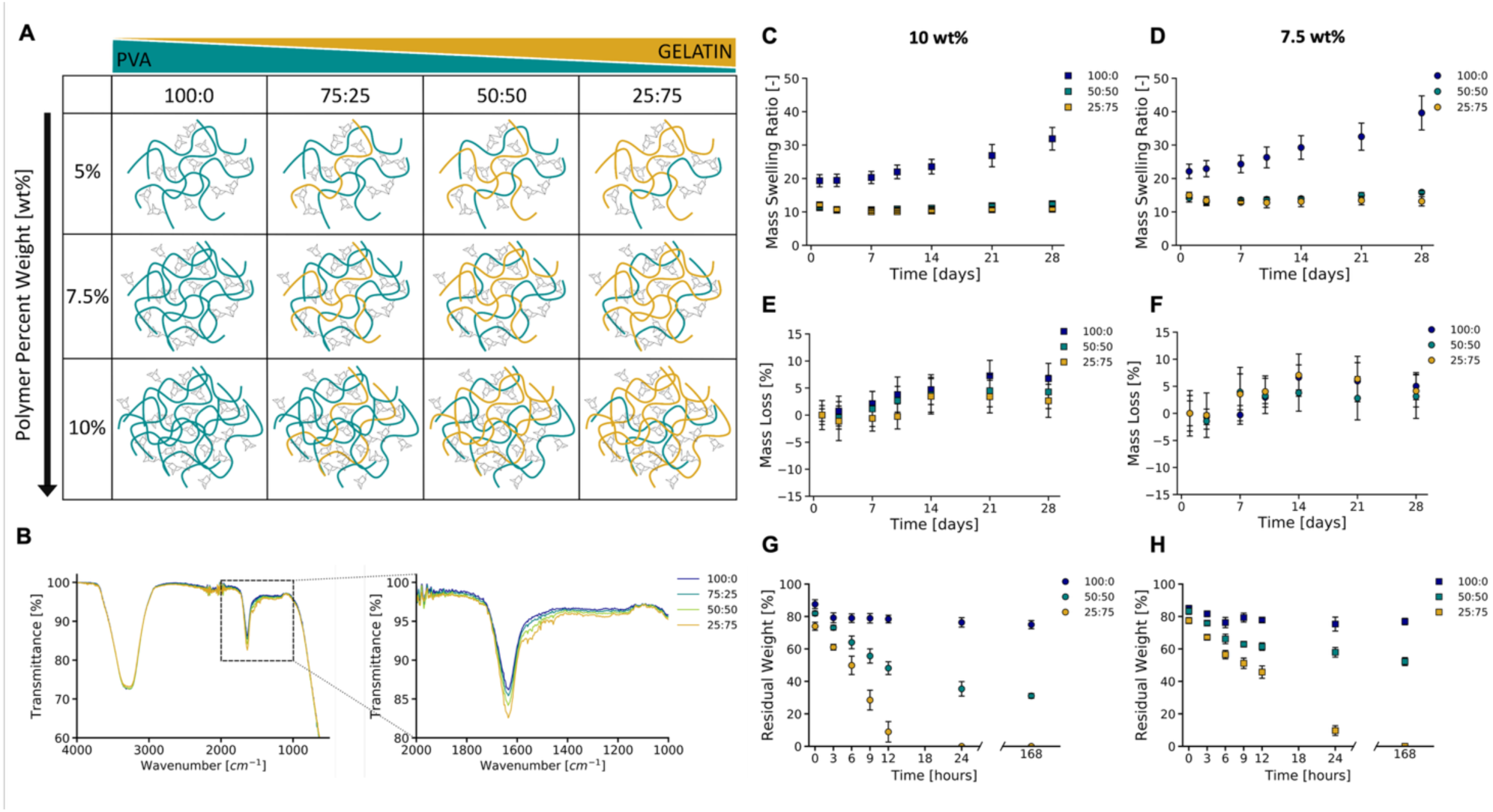
Physical properties and degradation profiles of the PVA-GEL hydrogel at different polymer percent weights (wt %) and polymer ratios (PVA:GEL). (A) Schematic showing the different formulations of PVA-GEL hydrogels tested. (B) FTIR spectra of 10 wt% PVA-GEL hydrogels at different polymer ratios after photopolymerisation. (C-D) Mass swelling ratio of 10 wt% and 7.5 wt% PVA-GEL hydrogels with different polymer ratios. (E-F) Changes in mass loss percentages in 10 wt% PVA-GEL hydrogels and 7.5 wt% PVA-GEL hydrogels with different polymer ratios after the initial soluble fraction (overall mass loss is reported in Figure S2). (G-H) Enzymatic degradation profiles of (G) 10 wt% and (H) 7.5 wt% PVA-GEL hydrogels with different polymer ratios after being immersed in a collagenase solution. All reported data represent the mean of 3 repeats (n=3), each of them consisting of at least 3 replicates (N=9). All results are expressed as the mean ± the standard deviation of the means. One-way ANOVA was performed to compare the means of the groups and a Tukey-Kramer test was used for multiple comparisons between the means. Differences were considered significant at a significance level of 5% (* p < 0.05), 1% (** p < 0.01) or 0.1% (*** p < 0.001). Statistical analyses results shown in Table S1, S2, S5, S6.

Hydrogel swelling behavior was evaluated by measuring the mass swelling ratio over 28 days to match the physiological timeline for neural tissue ingrowth^61^. Pure PVA hydrogels exhibited significant swelling, increasing up to 31.88 ± 3.39 for 10 wt% and 39.63 ± 5.10 for 7.5 wt% after 28 days (mean ± standard deviation, **Figure 3C-D**). In contrast, scaffolds with 50:50 or 25:75 PVA:GEL ratios maintained stable swelling profiles over time. A similar trend was observed for both polymer weight percentages investigated (i.e., 10 wt% and 7.5 wt%). This reduction in water uptake could be attributed to increased crosslinking density and lower hydrophilicity caused by the substitution of the hydroxyl groups in PVA with NB and the higher DS of GEL-NB^62^. As lower swelling ratio is ideal to minimise unwanted pressure at the implant site, the introduction of GEL in PVA hydrogels introduces a clear benefit for neural interfaces with the constrained CNS tissues.

Biodegradation kinetics were investigated by assessing hydrolytic mass loss in DPBS and enzymatic degradation in a collagenase solution. Mass loss was investigated to gain insight into the rate of intrinsic hydrolytic degradation of PVA-GEL hydrogels over time, which affects the stability of the implant. Mass loss profiles showed an initial burst release of 30 % soluble content between day 0 and day 1, due primarily to unreacted polymer (**Figure S2**). After normalizing by the soluble fraction, which corresponds to the percentage of polymer released after the first 24 hours of incubation (**Figure S2**), a consistent increase in mass loss was observed between day 3 and day 14 (**Figure 3E-F**). No significant changes were observed from day 21 to day 28, similarly to other hydrogel systems^46,63^. This is a desirable behavior to provide initial structural persistence and gradual ECM turnover following implantation. PVA-GEL hydrogels showed an increase in soluble fraction with increasing gelatin content, owing to the presence of larger amounts of unreacted GEL-NB polymer released during the first hours of incubation (**Figure S2**). This behavior could be caused by the higher steric hindrance of gelatin, that can decrease crosslinking efficiency and lead to less tightly crosslinked hydrogels^64,65^. Enzymatic degradation via a collagenase assay was performed to evaluate scaffold biodegradability and to confirm that chemical functionalization and photo-polymerisation did not alter this property^50,66^. GEL-containing scaffolds were susceptible to collagenase-mediated breakdown, with steady rates of degradation over 7 days of incubation increasing alongside GEL content (**Figure 3 G-H**). As anticipated, faster degradation rates were observed for 7.5 wt% hydrogels compared to their counterparts at 10 wt% due to their lower crosslinking density^66^. After 7 days of incubation in collagenase solution, 10 wt% and 7.5 wt% PVA-GEL hydrogels at a 50:50 (PVA:GEL) polymer ratio reached a residual weight of 52.24 ± 2.44 % and 31.10 ± 1.49 %, respectively. The fastest degradation rate was observed for PVA-GEL hydrogels with the highest concentration of gelatin (25:75, PVA:GEL) that were completely degraded after 24 hours (7.5 wt% formulation) and 7 days (10 wt% formulation). These results confirmed that NB functionalization and photo-crosslinking did not impair the intrinsic degradability of GEL, which remained accessible to cell-secreted proteases that mediate physiological tissue remodelling.

Biomaterial stiffness influences cell behaviour and glial scar encapsulation in brain implants. Mechanical properties were assessed via unconfined compression to determine the Young’s modulus of hydrogels at different timepoints following incubation in DPBS (i.e., day 0, 7, 14, 21, and 28). Initial moduli for 10 wt% formulations ranged from 12.34 ± 2.69 kPa (100:0) to 5.51 ± 0.84 kPa (25:75), decreasing over time due to swelling and degradation of the scaffolds (**Figure 4A-B**). After 28 days of incubation, the Young’s moduli significantly decreased to 1.48 ± 0.59 kPa (100:0) and 2.92 ± 1.06 kPa (25:75). These values fall within the physiologically relevant range for scaffolds designed to mimic the properties of the brain^67,69^, and the observed softening over time reflected the gradual relaxation and degradation expected *in vivo*. Lower polymer content (7.5 wt%) resulted in softer scaffolds across all composition, in line with previous studies^30^. Notably, the 25:75 PVA-GEL hydrogels maintained a modulus near 3 kPa after 28 days, which closely approximates the stiffness of healthy cortical tissue (∼1–3 kPa)^68^.

**Figure 4.**
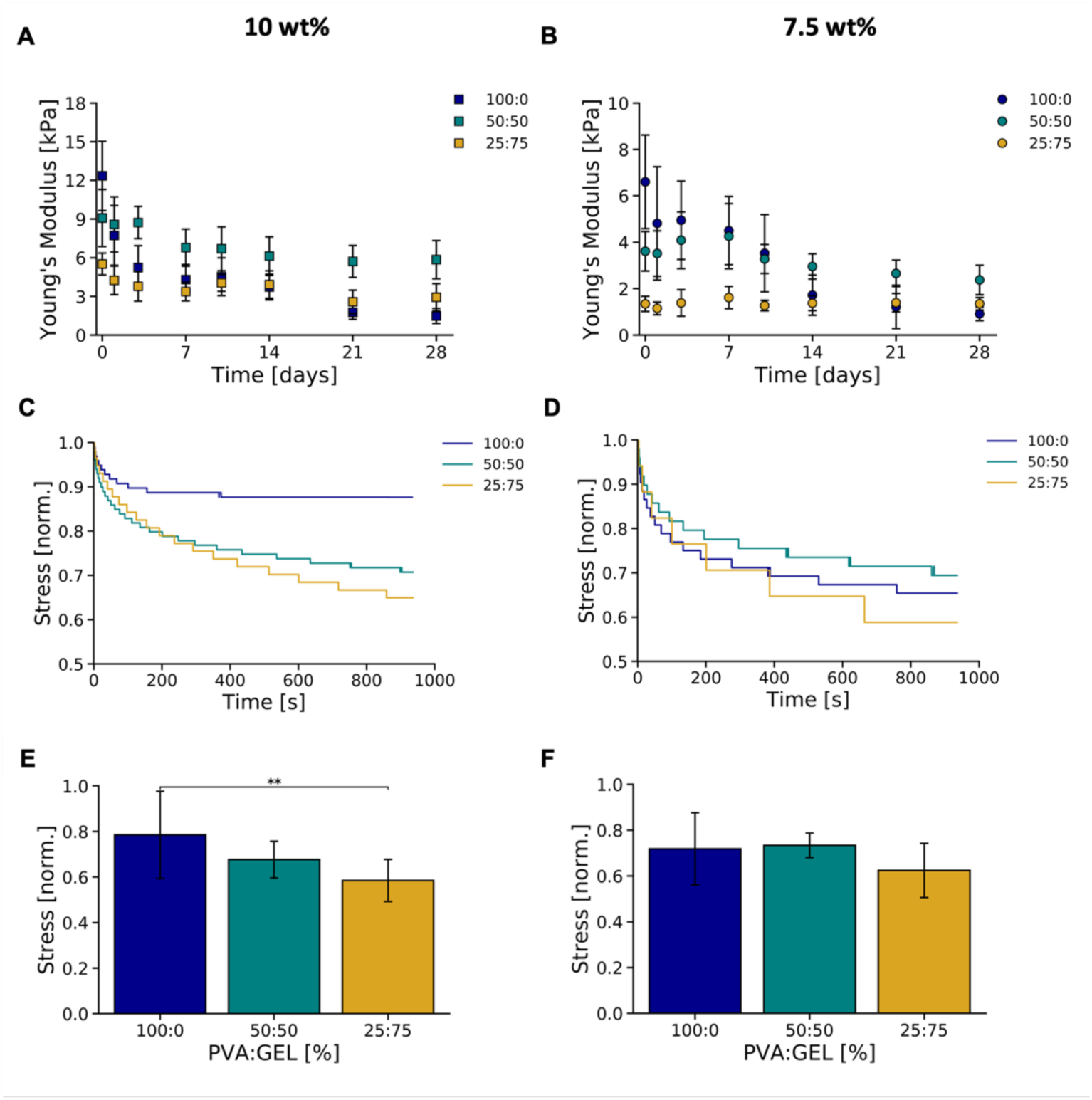
Mechanical properties of the PVA-GEL hydrogel at different polymer percent weights (wt %) and polymer ratios (PVA:GEL). (A-B) Young’s moduli of 10 wt% and 7.5 wt% PVA-GEL hydrogels with different polymer ratios. (C-D) Representative stress-relaxation curves of 10 wt% and 7.5 wt% PVA-GEL hydrogels after 7 days of incubation in DPBS. (E-F) Stress-relaxation values of 10 wt% and 7.5 wt% PVA-GEL hydrogels reached after 15 min of constant strain. All reported data represent the mean of 3 repeats (n=3), each of them consisting of at least 3 replicates (N=9). All results are expressed as the mean ± the standard deviation of the means. One-way ANOVA was performed to compare the means of the groups and a Tukey-Kramer test was used for multiple comparisons between the means. Differences were considered significant at a significance level of 5% (* p < 0.05), 1% (** p < 0.01) or 0.1% (*** p < 0.001). Statistical analyses results shown in Table S7, S8.

The viscoelastic behavior of PVA-GEL hydrogels was assessed, reflecting this emerging design criterion for scaffolds used for neural tissue engineering^70^. Viscoelasticity is particularly relevant to brain interfacing applications, as neural tissues are among the most viscoelastic structures in the body^20^. However, covalently crosslinked hydrogels often exhibit a fully elastic behavior, which has been shown to hinder cell development, growth, and migration^58,71,72^. Stress relaxation was measured after 7 days of swelling in DPBS under a constant strain of 15% compression for 15 minutes. These results showed that PVA-GEL scaffolds dissipated 10-40% of the applied stress, with a general trend of 25:75 PVA-GEL being more viscoelastic than pure PVA hydrogels (**Figure 4E-F**, stress values after 24 h are reported in **Figure S2**). Although this behavior was similar to other PVA-and GEL-based hydrogels reported previously in the literature^73,74^, the native viscoelastic properties of the brain are significantly different^75^. Overall, increasing GEL content was shown to enhance viscoelastic behavior, which was likely due to the higher crosslinking density and the composition of this biopolymer. However, no consistent correlation was observed between polymer ratio, weight percentage, and relaxation behavior across all timepoints, which suggested that other factors such as crosslinking topology and local network heterogeneity played a more dominant role. Moreover, these findings also aligned with recent studies showing that viscoelastic properties are governed more by crosslinking chemistry and network dynamics than by the nature of the base polymer alone^57^.

These results demonstrated that the physical properties of PVA-GEL hydrogels could be finely tuned by varying the ratio of PVA to GEL and the total polymer concentration. In particular, the 10 wt% and 7.5 wt% formulations with a 25:75 PVA:GEL ratio exhibited low swelling, consistent biodegradability, as well as soft yet stable mechanical properties. Moreover, the tailorability of the PVA-GEL system enabled the systematic design of scaffolds for neural interfaces that could support the growth and development of robust astrocytic constructs.

### 2.3 Engineering astrocyte-supportive microenvironments with PVA-GEL hydrogels

Astrocytes play a central role in neural development and homeostasis by providing trophic support, modulating synaptic activity, and orchestrating ECM remodelling^76–80^. Astrocytic cells have also been shown to respond to spatial and topological cues via mechanical activation, resulting in modulation of cell migration and network development^81,82^. Therefore, engineering defined microenvironments that support astrocyte growth and function constitutes an attractive strategy for the fabrication of biomimetic neural constructs. Although previous studies have investigated the interactions between astrocytes and neurons *in vitro*^38,78,83–88^, a systematic investigation of the optimal conditions required for the culture of primary astrocytes in 3D systems remains largely unexplored. Following the identification of PVA-GEL hydrogel formulations with desirable physical and mechanical properties, they were evaluated for their ability to sustain astrocyte growth and promote key hallmarks of glial function, including matrix remodelling and cell adhesion.

Primary astrocytes isolated from postnatal day 4 (P4) rats were encapsulated in PVA-GEL hydrogels with varying polymer content (7.5 wt% and 10 wt%) and PVA:GEL ratios (100:0, 75:25, 50:50, 25:75) and maintained in culture for 7 days *in vitro* (DIV). To identify optimal material compositions for cell growth, motility and ECM degradation capabilities, immunofluorescent staining (IFS) was carried out against glial fibrillary acidic protein (GFAP), paxillin, and MMP-2. From qualitative observations of cell morphology, IFS revealed that astrocytes in hydrogels with high gelatin (25:75 PVA:GEL) content exhibited elongated morphologies and extended cellular processes, while those in pure PVA hydrogels (100:0, PVA:GEL) showed round and restricted morphologies at all polymer contents (**Figure 5**). Hydrogels with 5 wt% polymer content were included in the investigation to evaluate astrocytic growth on softer substrates. 5 wt% hydrogels with medium to high gelatin content (50:50 and 25:75 PVA:GEL) degraded completely after 3-4 days in culture. This could be explained in part due to the high porosity of hydrogels with low polymer content, which provides the cells with more space to develop and remodel the scaffold.

**Figure 5.**
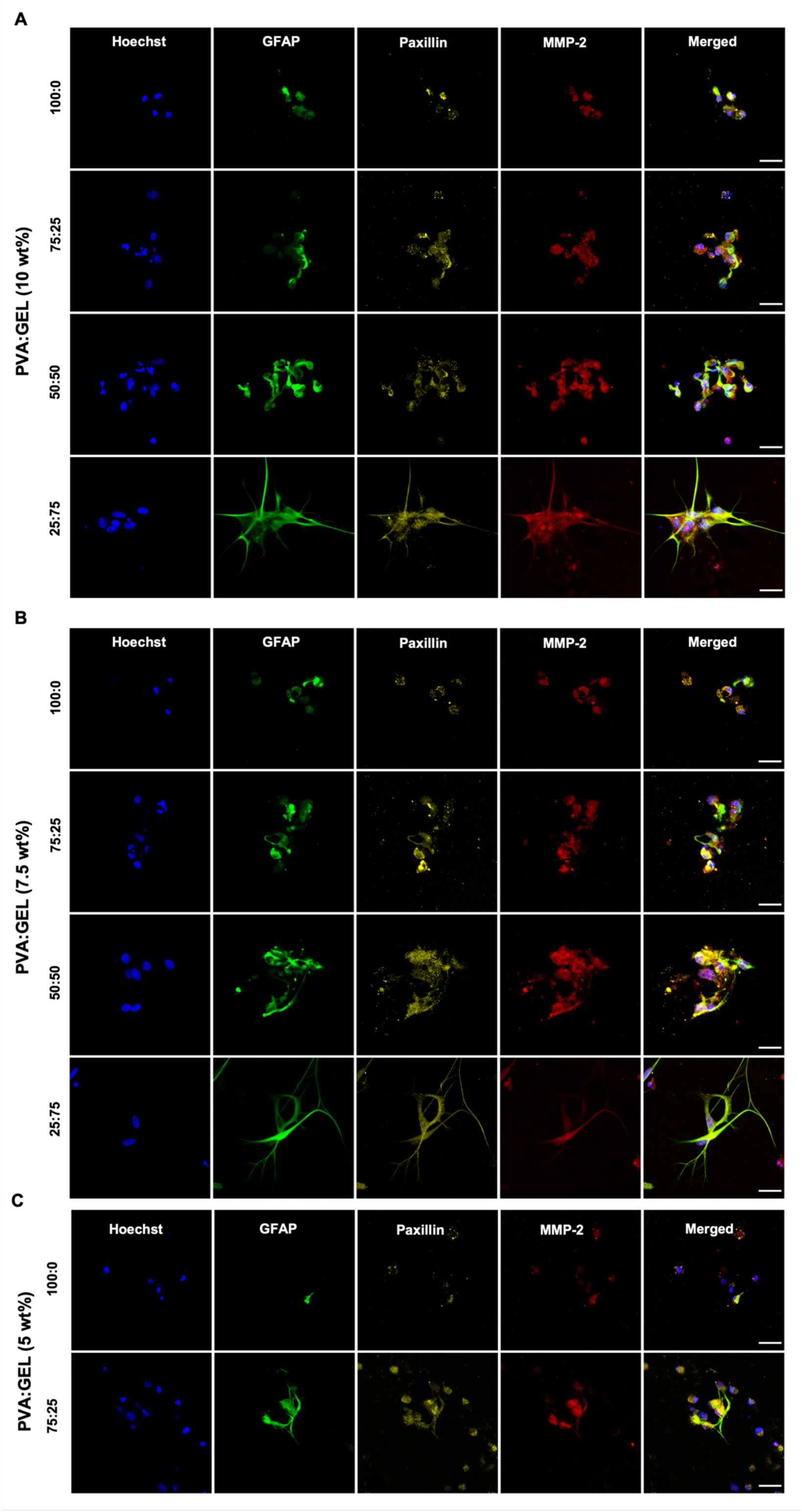
Representative immunofluorescent micrographs of primary astrocytes encapsulated PVA-GEL hydrogels at DIV7. (A) 10 wt%, (B) 7.5 wt%, (C) 5 wt% PVA-GEL. Cell nuclei are shown in blue (Hoechst), the expression of GFAP in green, paxillin in yellow, and MMP-2 in red. Scale bar = 30 µm.

Cell viability was assessed using a resazurin-based metabolic assay at DIV7 post-encapsulation. Astrocytes grown in gelatin-containing hydrogels exhibited significantly higher rates of metabolic activity compared to those in pure PVA formulations (**Figure 6A**). Among all conditions tested, astrocytes encapsulated in 10 wt% PVA-GEL hydrogels with a 25:75 ratio exhibited the highest levels of cell viability at day 7 (p < 0.01). These results were consistent with previous reports showcasing the high cytocompatibility of gelatin-containing scaffolds, attributed to the provision of critical adhesion sites and pro-survival cues^89^. Here, the inclusion of gelatin mitigated the loss in viability that was observed in previous living electrode constructs^11,41^ due to dense hydrogel networks with high crosslinking density. These findings suggest that the biochemical composition of the matrix and the inclusion of adhesion-promoting polymers, can effectively offset the limitations imposed by higher crosslinking densities. Quantitative analysis of fluorescence micrographs further confirmed that astrocytes in 25:75 PVA-GEL hydrogels displayed significantly greater GFAP expression and cell elongation compared to other formulations (**Figure 6B–D**). These findings indicated that the combined biochemical and biophysical properties of this formulation synergistically promoted glial process elaboration, which is a key indicator of astrocyte maturity and function^85^.

**Figure 6.**
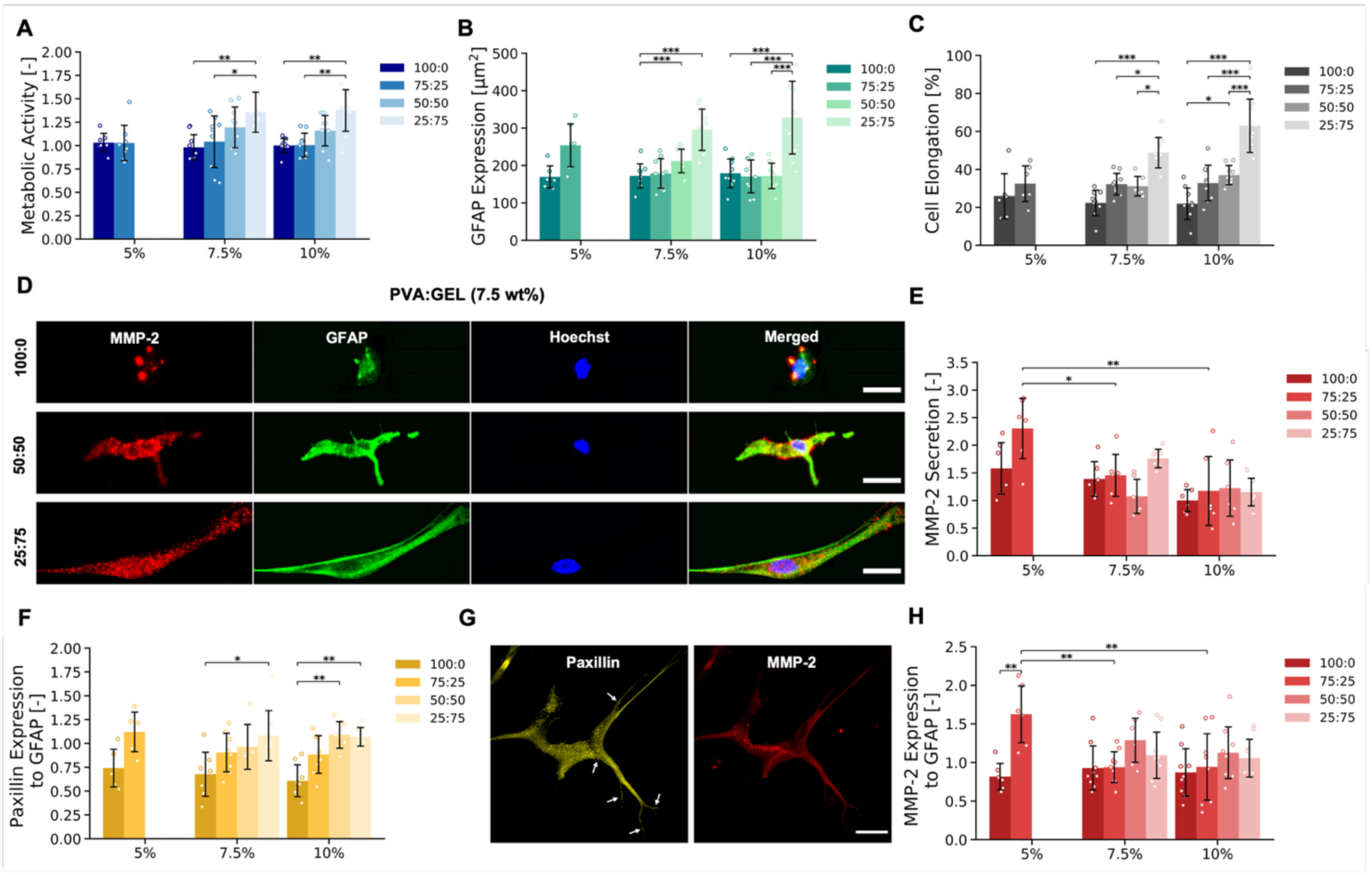
Cell viability and adhesion in primary astrocytes encapsulated in PVA-GEL across polymer percent weights and polymer ratios at DIV7. Primary astrocytes were encapsulated for 7 days in PVA-GEL hydrogels upon fixation. (A) Metabolic activity measured with resazurin-based assay. (B) GFAP expression per cell. (C) Percentage of elongated cells quantified from GFAP morphology. (D) Representative immunofluorescence micrographs of primary astrocytes encapsulated in 7.5 wt% PVA-GEL hydrogels. Cell nuclei are shown in blue, GFAP expression in green, MMP-2 in red. A concomitant increase in cell elongation was observed with the increase of gelatin content in the hydrogels. Scale bar = 20 µm. (E) MMP-2 secretion quantified via zymography. (F) Paxillin and (H) MMP-2 expression relative to the expression of GFAP (G) Representative immunoflurescence micrographs showing paxillin and MMP-2 expression in primary astrocytes encapsulated in 10 wt% PVA-GEL hydrogels. Paxillin is shown in yellow, and MMP-2 in red. Paxillin structures denoting the formation of mature focal adhesions are indicated by white arrows. Scale bar = 30 µm. All reported data represent the mean of 3 repeats (n=3), each consisting of at least 3 replicates (N=9). All results are expressed as the mean ± the standard deviation of the means. One-way ANOVA was performed to compare the means of the groups and a Tukey-Kramer test was used for multiple comparisons between the means. Differences were considered significant at a significance level of 5% (* p < 0.05), 1% (** p < 0.01) or 0.1% (*** p < 0.001).

The cell adhesion and ECM remodelling behaviour of astrocytes in the hydrogel was further evaluated using gel zymography to quantify levels of secreted MMP-2, which is an important mediator of ECM turnover in the CNS^94^. MMP-2 expression plays an important role in astrocyte morphology, migration, synaptogenesis and synaptic plasticity in the adult CNS^95^. These results showed that astrocytes in gelatin-rich hydrogels were shown to secrete significantly more active MMP-2 than those with higher PVA content (**Figure 6E**). However, no significant differences in MMP-2 secretion were observed between hydrogels with different PVA:GEL ratios at both 7.5 wt% and 10 wt%. Quantitative assessment of fluorescence micrographs showed that paxillin expression, a biomarker involved in the formation of focal adhesions, increased concomitantly with the increase of gelatin content (**Figure 6F-G**). Focal adhesions are key mediators of astrocyte morphology, growth, and migration *in vitro*, and are involved in the regulation of neural synapses and neurotransmitter release^58^. Quantitative assessment of MMP-2 expression also revealed increased protease content in gelatin-rich hydrogels, with no significant differences between hydrogels with different PVA:GEL ratios and polymer contents (**Figure 6G-H**). In line with the GFAP expression results, the high crosslinking density of hydrogels with low polymer content may have allowed the cells to detect more gelatin adhesion cues and produce higher levels of MMP-2. However, further investigations are required, material stiffness could also have influenced protease activity^96^.

In summary, these results demonstrated that PVA-GEL hydrogels, particularly those formulated with 25:75 PVA:GEL at 10 wt%, could provide a permissive microenvironment for astrocyte survival, maturation, and ECM remodelling. Based on the physical and mechanical characterization of the scaffolds, primary astrocytes seemed to be best supported by lower Young’s moduli (1-6 kPa, **Figure 4A-B**), lower mass-swelling ratios (**Figure 3C-D**), and more easily degradable hydrogels (**Figure 3G-H**). The formulation with 10 wt% polymer content and 25:75 PVA-GEL ratio was selected for subsequent testing because of the higher stiffness compared to the 7.5 wt% formulation, as higher Young’s moduli are expected to improve the stability and ease of implantation of living electrode devices. It was hypothesised that these PVA-GEL hydrogels could provide a robust foundation to engineer complex neural constructs by supporting the co-development of astrocytes and NPCs into synaptically active neural constructs.

### 2.4 Astrocyte-driven maturation of neural constructs

After the *in vitro* tests showed that scaffolds with 25:75 PVA:GEL polymer ratio and 10 wt% polymer content provided a supportive environment for astrocyte growth, the formulation was used to investigate whether the material could support the development of neuro-glia constructs. This approach, entailing co-cultures of astrocytes and NPC-enriched neurospheres, aimed to show that physiological astrocyte-mediated cues^97^ can be leveraged to regulate NPC fate, neuronal differentiation, and synaptic maturation in tissue engineered constructs.

To assess the contribution of astrocytes to neural network development, constructs of NPC-enriched neurospheres were encapsulated with and without P4 hippocampal astrocytes. NPCs were first isolated from the ventral mesencephalon of day 14 (E14) rat embryos and grown as neurospheres in ultra-low attachment culture substrates for DIV7-10 at an average diameter of 50-100 µm. NPC-enriched neurospheres were then encapsulated in 10 wt% PVA-GEL hydrogels (25:75) either with or without astrocytes, and maintained *in vitro* to allow for construct maturation for either DIV7 or DIV14. At these timepoints, metabolic activity was assessed using a commercial resazurin-based assay (AlamarBlue), and constructs were then processed for IFS against GFAP for astrocyted, βIII-tubulin for neurons, myelin, synaptophysin, and PSD-95. IFS analysis revealed striking differences in construct composition and morphology between the two conditions. From qualitative observations, co-cultures exhibited extensive neurite outgrowth with elongated axonal tracts traversing large regions of the scaffolds, while neurosphere-only constructs showed reduced and network formation overall (**Figure 7A-B,G-H**). Constructs at DIV14 exhibited consistently higher differentiation and organisation compared to those at DIV7, which was indicative of progressive maturation over time (**Figure 7A-B**). Widespread expression of the neuronal marker βIII-tubulin was observed in DIV14 co-cultures, which confirmed successful neuronal differentiation from encapsulated progenitor populations.

**Figure 7.**
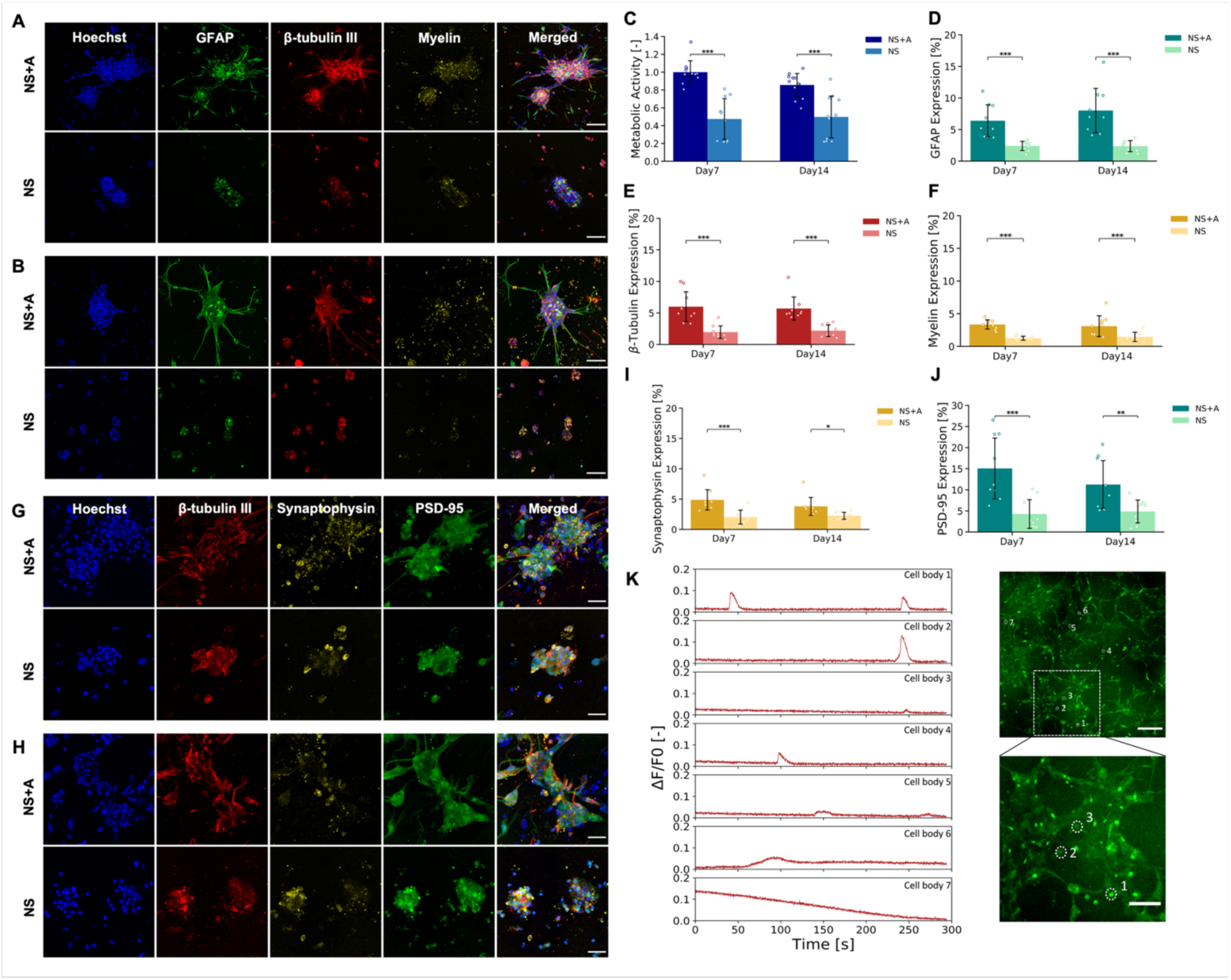
The influence of astrocytes on viability, cell identity and functionality in the PVA-GEL 3D co-cultures over time. (A-B) Representative fluorescent micrographs of primary neurospheres encapsulated in 10 wt% 25:75 PVA-GEL hydrogels with and without P4 astrocytes at DIV7 (A) and DIV14 (B). Cell nuclei are shown in blue (Hoechst), the expression of GFAP in green, βIII tubulin in red, myelin in yellow. Scale bar = 100 µm. (C) Metabolic activity measured with resazurin-based assay, (D) GFAP expression, (E) βIII tubulin expression, (F) myelin expression, quantified in percentage of the image covered by the biomarker after thresholding the intensity to a pre-set value. (G-H) Representative fluorescent micrographs of primary neurospheres encapsulated in 10 wt% 25:75 PVA-GEL hydrogels with and without P4 astrocytes at DIV7 (G) and DIV14 (H). Cell nuclei are shown in blue (Hoechst), the expression of βIII tubulin in red, synaptophysin in yellow, PSD-95 in green. Scale bar = 100 µm. (I) Synaptophysin expression and (J) PSD-95 expression, quantified in percentage of the image covered by the biomarker after thresholding the intensity to a pre-set value. (K) Functional characterization of neural network activity in PVA-GEL hydrogel at DIV7. Calcium activity was recorded from co-cultures of neurospheres and astrocytes encapsulated in PVA-GEL hydrogels at DIV7. Representative ROIs containing the recorded cells surrounded by white dotted lines are shown in the immunofluorescence panel. Scale bar = 200 µm. All reported data represent the mean of 3 repeats (n=3), each of them consisting of at least 3 replicates (N=9). All results are expressed as the mean ± the standard deviation of the means. One-way ANOVA was performed to compare the means of the groups and a Tukey-Kramer test was used for multiple comparisons between the means. Differences were considered significant at a significance level of 5% (* p < 0.05), 1% (** p < 0.01) or 0.1% (*** p < 0.001).

Metabolic activity was significantly higher in the co-cultures compared to neurosphere-only constructs at both DIV7 and DIV14 (p < 0.01, **Figure 7C**). Quantitative image analysis further confirmed that co-cultures exhibited significantly higher expression levels of βIII-tubulin (**Figure 7D**) and GFAP (**Figure 7E**) compared to neurosphere-only constructs at both timepoints. The percentage of astrocytes and neurons within the hydrogels did not significantly vary across the different experimental conditions and timepoints tested (**Figure S3**), remaining similar to the 50/50 physiological ratio observed in the brains of rodents and humans^98^. These results also revealed enhanced myelination in the co-cultures at DIV7 and DIV14 (**Figure 7F**), which suggested enhanced maturation of oligodendrocyte lineage cells and enhanced functionality of the neural network. Synaptic maturation within the constructs was assessed via IFS against synaptophysin and PSD95, which are markers of pre-and post-synaptic specialisations, respectively. Increased expression of these markers was observed throughout the co-cultures, which suggested improved formation of putative synaptic contacts (**Figure 7G-H**). Quantification of synaptophysin and PSD95 density across the constructs indicated a high prevalence of synaptically active domains within the constructs in both timepoints (**Figure 7I-J**). To further validate the emergence of functional connectivity in encapsulated co-cultures, spontaneous calcium transients were recorded using Fluo-4 calcium imaging at DIV7 (**Figure 7K**) and DIV14 (**Figure S3**). Consistent Ca^2+^ signaling live-cell activity can be observed in the co-culture at both timepoints, strengthening the evidence on neural network development and functionality. These findings underscore the crucial role of astrocytes in establishing local microenvironments that support multi-lineage differentiation, structural organisation, and neural network formation *in vitro*^76,78^.

The concentration of brain-derived neurotrophic factor (BDNF) in the medium was also assessed as a proxy for the neurotrophic support of astrocytes, and the development of the neuronal population in the cultures (**Figure S3)**. The role of D-Serine, a gliotransmitter secreted by astrocytes, has also been assessed across conditions (**Figure S3)** to gain insight into the potential mechanism underlying glial support. D-serine has been shown to increase at early developmental stages of the mammalian brain, and it is known to be involved in neural circuit refinement^45,99^. However, no detectable levels of BDNF or D-and L-Serine could be observed with ELISA assays, across conditions and timepoints (**Figure S3**). Similar results have been reported for other hydrogel-based systems, where protein retention, adsorption, and limited mass transport obscured soluble factor detection in bulk media^100,101^. Therefore, these factors may still play localized roles within the scaffold microenvironment, and more sensitive or spatially resolved detection methods are required to gain further insight into their involvement in construct maturation.

Overall, this work demonstrates that astrocyte-enriched PVA-GEL scaffolds effectively support the emergence of synaptically competent neural networks from NPC-enriched neurospheres. This straightforward approach bypassed the need for pre-differentiated or spatially patterned cells, by leveraging endogenous neuro-glia interaction mechanisms. These biomimetic constructs hold broad applicability for emerging applications in the development of tissue-engineered coatings for in biohybrid neural interfaces, as well as in fundamental research and translational applications.

### 2.5 *Ex Vivo* integration of neural constructs with organotypic brain slices

Lastly, an *ex vivo* model of organotypic hippocampal slices was used to study tissue-level integration, in a controlled environment that closely replicates the cytoarchitecture and multicellular dynamics of the native brain. This system was used to validate the suitability of the engineered constructs as a substrate for neural tissue ingrowth and for supporting synaptic connectivity between encapsulated cells and host neural tissue.

Organotypic hippocampal slices were generated from P4 rats and grown on an air-liquid interface^102^ to study the PVA-GEL tissue interface. Culture conditions were first optimized to ensure preservation of tissue viability and cytoarchitecture of slices grown on pristine PVA-GEL scaffolds (**Figure S4**). Organotypic cultures were then formed on top of pristine (PVA-GEL only) hydrogels to evaluate the ability of the scaffold to support the attachment and ingrowth of physiological neural tissue (**Figure 8A**). From qualitative observations, phase-contrast and fluorescence imaging showed adhesion to the scaffold after DIV21 in culture (**Figure 8B-C**). GFAP^+^ astrocytes seemed to migrate out of the slices and into the scaffold, while forming membrane projections with paxillin+ focal adhesions (**Figure 8C**). This underscored the biocompatibility of the PVA-GEL system and its potential to support the formation of stable and structurally integrated interfaces with host tissues.

**Figure 8.**
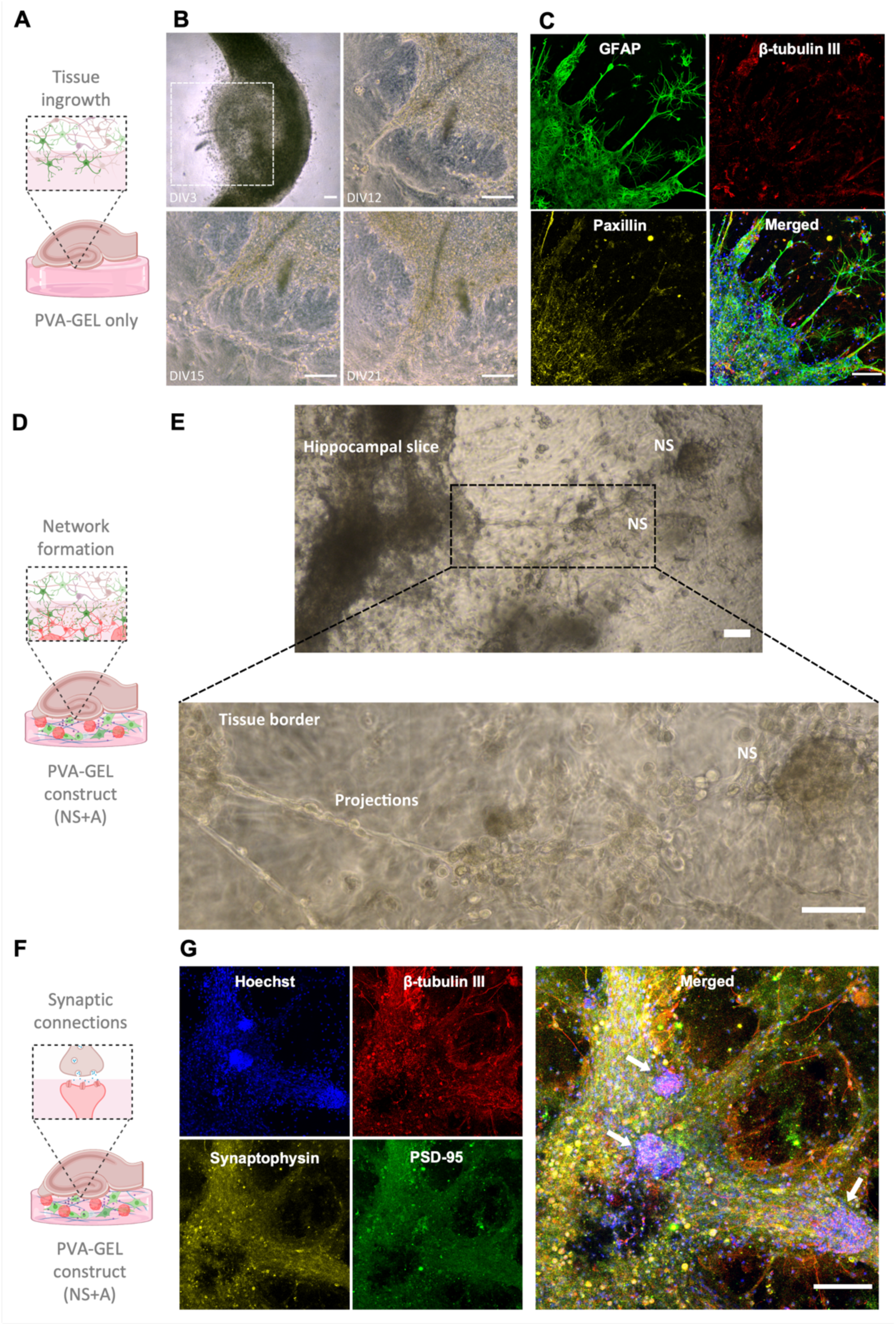
Evaluation on the interface properties of the PVA-GEL in chronic organotypic cultures. (A) Schematic of the PVA-GEL-only characterization, where tissue ingrowth from an organotypic hippocampal slice grown on an air-liquid interface is observed over time. (B) Representative phase-contrast micrographs of the tissue edge of the organotypic hippocampal slices cultured on PVA-GEL hydrogels over DIV21. Images were acquired at DIV3, DIV12, DIV15 and DIV21. The dotted-line area at DIV3 shows the area captured in the following images. Scale bar = 150 µm. (C) Representative fluorescent micrographs of organotypic hippocampal slice cultures on PVA-GEL hydrogels at DIV21. Cell nuclei are shown in blue (Hoechst), the expression of GFAP in green, βIII tubulin in red, paxillin in yellow. Scale bar = 100 µm. (D) Schematic of the PVA-GEL-only characterization. (E) Representative phase-contrast micrographs of hippocampal slices interfacing with neurospheres encapsulated in a PVA-GEL hydrogel at DIV21. Scale bar = 100 µm. (F) Schematic of the PVA-GEL-only characterization. (G) Representative immunofluorescence micrographs of organotypic hippocampal slices cultured on top of PVA-GEL constructs at DIV14. Cell nuclei are shown in blue, expression of βIII tubulin in red, synaptophysin in yellow, and PSD-95 in green. All fluorescent channels merged are presented and neurospheres are indicated by white arrows. Scale bar = 100 µm.

Organotypic brain slices were then formed on top of co-cultures encapsulated in the hydrogels to evaluate neural network formation (**Figure 8D**) and synaptic integration (**Figure 8F**) of the living electrode concept with physiological neural tissue. Hippocampal cultures could effectively be formed and maintained in direct contact with the neural construct for up to DIV21, developing migratory edges and extensive tissue projections that colocalized with NPC-enriched neurospheres encapsulated within the scaffold (**Figure 8E**). Fluorescent micrographs revealed widespread expression of the neuronal marker βIII-tubulin within the slice. Both pre- and post-synaptic markers were expressed across the construct (**Figure 8G**). These results present an early indication of the preservation of the native cytoarchitecture in organotypic slices, as well as the formation of putative synaptic contacts with the underlying construct.

This organotypic model provided a versatile platform to investigate the early phases of construct– tissue integration in a high-resolution and ethically tractable system. However, the lack of fluorescent reporters that could distinguish between neural cells from the construct and those from the organotypic slice precluded the ability to analyse the behavior of both populations independently. Therefore, future studies will incorporate selective labelling strategies and functional imaging tools to investigate the origin and electrophysiological maturity of synaptic contacts. Despite this limitation, the ability of the PVA-GEL system to support tissue interfacing *ex vivo* underpins its potential to foster construct maturation and biological integration *in vivo*.

## 3. Conclusion

By enhancing a versatile biomaterial system with a biomimetic strategy, we provide a straightforward method to produce robust and synaptically competent neural constructs with broad applicability. This approach harnesses endogenous astrocytie-guided mechanisms of neural network development to drive cell differentiation and network maturation, instead of relying on patterning techniques or pre-differentiated phenotypes.

The hydrogel system is based on NB-functionalized PVA and gelatin, crosslinked via visible light-triggered thiol–ene chemistry to yield a cytocompatible and highly tuneable matrix. A systematic optimization of the hydrogel formulation yielded scaffolds that effectively supported astrocyte growth and development, while providing programmable structural integrity and biodegradability. A 10 wt% nominal polymer content and 25:75 PVA-GEL ratio was identified as the optimal PVA-GEL hydrogel composition to promote the growth and development of primary astrocytes, which highlighted the key role of bioactive and mechanical cues in the modulation of glial cell behaviour. These scaffolds were shown to support multi-lineage differentiation and self-organisation of encapsulated NPCs into robust neural constructs. In contrast, constructs formed without a glial component exhibited limited maturation, which underscores the central role of astrocytes in neuronal differentiation, neurite extension, and functional network maturation. *Ex vivo* culture with organotypic brain slices showed evidence of direct structural integration between PVA-GEL hydrogels and physiological neural tissue. Cellular processes extended across the tissue-construct interface, and the expression of pre-and post-synaptic markers suggested the development of synaptic networks. Although further experimental validation is needed in an *in vivo* environment, these findings suggest that this approach can yield mature neural constructs capable of integrating and communicating with physiological neural tissues.

PVA-GEL-based constructs not only recapitulate essential aspects of neural tissue organisation, but they also possess the functional and structural features necessary to bridge the interface between bionic implants and the nervous system. The strategy presented here departs from traditional biofabrication approaches, which often rely on patterning defined phenotypes into specific spatial arrangements. Instead, this method harnessed endogenous astrocytic pathways to produce biomimetic neural architectures. This bottom-up approach offers new opportunities for engineering physiologically relevant constructs with minimal processing, biological fidelity, and translational relevance.

Although recent studies have demonstrated the benefits of soft, stretchable electronics and tissue-permissive materials^101^, few platforms have focused on incorporating living neural cells to form functional synapses with the host^103^. This approach builds on the living electrode concept^11,41^ by introducing significant improvements in construct viability, scaffold design, and *ex vivo* validation. It has been shown that the multi-layered hydrogel coatings based on this PVA-GEL system could be effectively formed on platinum electrode substrates without compromising electrochemical performance or eliciting cytotoxic responses^43^. Future work will focus on the engineering of implantable microelectrode PVA-GEL coatings and *in vivo* validation of this new iteration of the living electrode technologies. Further improvements could be implemented in the hydrogel system, such as additional functionalizations of the NB groups, such as introducing topographical cues in the system to direct cell migration, or biological cues to direct cell fate. The introduction of electrical stimulation cues constitutes another avenue for improving the neural development and functionality of the construct^104,105^, leveraging the implanted stimulation device. Thus, the utility of this platform extends beyond neural interfacing, as the modularity and physiological relevance of the constructs make them ideally suited for applications in regenerative medicine, disease modelling, or drug screening.

Ultimately, this study advances the field of engineered neural interfaces by introducing a biomimetic framework for fabricating tissue-like neural constructs. This will in turn contribute to the realisation of next generation neurotechnologies that are no longer inert and foreign, but biologically integrated extensions of the nervous system itself.

## 4. Experimental Section

### Synthesis of Norbornene-Functionalised Polymers

#### PVA-NB Synthesis

PVA-NB synthesis was performed according to a protocol adapted from Qin *et al*^50^. Briefly, 1 g of PVA (13-23 kDa, 98% hydrolysed, Cat. #348406) and 2.3 mg of sodium p-toluenesulfonate (pTS) (Cat. #152536) were dissolved in 20 ml of anhydrous dimethyl sulfoxide (DMSO, VWR 23500.322, dried over 4-Å molecular sieves) for 1 hour at 60°C under nitrogen atmosphere. Separately, 0.3 g of cis-5-norbornene-endo-2,3-dicarboxylic anhydride (NB) (Cat. #247634) were dissolved in 5 ml of anhydrous DMSO at room temperature (RT) under nitrogen atmosphere. The NB solution was then added dropwise to the fully dissolved PVA solution, and the reaction was stirred for 16 hours at 50°C under nitrogen atmosphere. The product of the reaction was then dialysed (dialysis tubing cellulose membrane, 14 kDa cut-off, Cat. #D9527) for 24 hours against 100 mM sodium bicarbonate (NaHCO3, Cat. #S5761) in distilled water (DI), followed by 48 hours of dialysis in pure DI water (changed with fresh DI water at least three times per day). Lyophilisation was used to recover the chemical product and ^1^H-NMR was run to confirm the chemical reaction. The DS was determined by comparing the integral of the peak associated with the norbornene group (peak location δ=6.2) to the integral of the peak associated with protons on the PVA backbone (peak location δ=4). Reagents calculations were stoichiometrically performed to reach a DS equal to 7% (∼ 25.4 groups of norbornene per PVA chain, Figure S1).

#### GEL-NB Synthesis

GEL-NB synthesis was performed according to a protocol adapted from Koshy et al^30^. Briefly, 1 g of gelatin type A from porcine skin (Cat. #G1890) was dissolved in 100 ml of 0.1 M 2-(N -morpholino)ethanesulfonic acid (MES, Cat. #3671) at pH 6 and 37°C. Once the gelatin completely dissolved, 5-norbornene-2-methylamine (NB, TCI Cat. #N0907) was added to the reaction at a molar ratio of 3 mmol per gram of gelatin. NHS (Cat. #130672) and N-(3-Dimethylaminopropyl)-N′-ethylcarbodiimide hydrochloride (EDC, Cat. #03450) were then added to the reaction at a molar ratio of 1:3:1 (NB:EDC:NHS). The reaction was stirred for 4 hours at 37°C. The product of the reaction was then dialysed (dialysis tubing cellulose membrane, 14 kDa cut-off, Cat. #D9527) for 3 days against DI (changed with fresh DI water at least three times per day). ^1^H-NMR was run to confirm the chemical reaction. The number of moles of NB conjugated to the gelatin chain were determined by comparing the integral of the peak associated with the norbornene group (peaks location δ=6.0-6.3) to the integral of the peak associated with the phenol groups on the gelatin backbone (peak location δ=7.3, Figure S1). The DS was calculated by comparing the number of moles of norbornene crosslinked to the gelatin backbone to the number of moles of carboxyl groups (COOH) present in the gelatin molecule (0.1267 mol/100 g of gelatin) The targeted DS was equal to 18% and the obtained value was 18.9% (Figure S1).

#### PVA-GEL Hydrogel Fabrication

PVA-NB and GEL-NB polymers were dissolved in DPBS at 37°C at different ratios, namely 100:0, 75:25, 50:50, 25:75, and 0:100 (PVA:GEL). The crosslinker solution was prepared by dissolving dithiothreitol (DTT, Thermo Fisher, #R0861) in DPBS at a thiol-to-ene stoichiometric ratio of 5:10, taking as reference the DS of the PVA-NB. Upon complete dissolution, the macromer solution was allowed to cool down to RT and the crosslinker solution was added. Lastly, the precursor solution was mixed with the initiator solution (Eosin Y disodium salt solution, #E6003, dissolved in DPBS) to reach a final concentration of 0.1 mM Eosin Y. The final volume of DPBS was calculated to reach a nominal polymer content of 5 wt%, 7.5 wt% or 10 wt% macromer concentration, depending on the chosen formulation. The prepared solution was then poured in PDMS disk moulds (10 mm diameter and 0.8 mm thickness for mechanical and swelling tests, 6 mm diameter and 0.5 mm thickness for cell encapsulation studies) and samples were exposed for 3 min to visible light (SOLIS-4C, Thorlabs) at 15mW/cm^2^ measured at 515 nm.

#### FTIR

FTIR measurements were performed using a Thermo Scientific Nicolet iS50 FTIR instrument. Three independent samples were tested for each hydrogel formulation, and each spectrum measurement is the average of 6 spectra.

### Physical and Mechanical Characterisation

#### Swelling and Mass Loss Characterisation

The physical characterization of the hydrogels was performed as described by Aregueta-Robles et al^46^. Swelling and mass loss studies were performed at the following timepoints: 0, 1, 3, 7, 10, 14, 21, and 28 days after photopolymerization. Three samples were prepared for each timepoint and the experiments were repeated three times (n=3, N=9). Immediately after polymerisation, hydrogels were weighed to determine their initial wet mass (m_i_). Three samples (day 0) were lyophilised to obtain their dry mass (m_d0_) and the effective macromer at day 0 was calculated using the following formula (**Equation 1**):

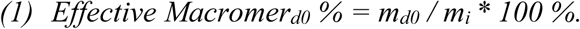

Each of the remaining samples was immersed in 3 ml of DPBS and incubated at 37°C until the defined timepoint was reached. Samples were then removed from the DPBS, blotted dry with wipes, and weighed to obtain their swollen mass (m_s_). All the samples were then lyophilised and weighed to determine their final dry mass (m_d_).

The effective macromer after day 0, was calculated as shown in **Equation 2**:

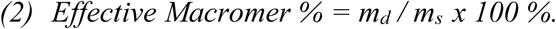

The mass swelling ratio (q), which indicates the increase in weight due to the water absorption under equilibrium conditions, was calculated as shown in eq 3:

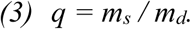

To calculate the mass loss, the initial dry weight (m_di_) of each sample was estimated, as shown in eq 4, using the effective macromer at DIV0:

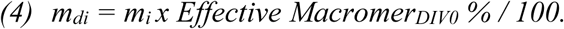

The mass loss, which indicates the amount of polymer that left the PVA-GEL network, was calculated as follows (**Equation 5**):

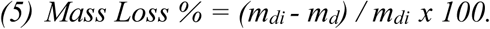

#### Compression Modulus

Uniaxial compression tests were performed to determine the compression moduli of PVA-GEL hydrogels. The bulk mechanical properties of the hydrogels were measured by uniaxial compression testing using a Bose ElectroForce 3200 test instrument. Compression tests were performed in the same conditions and at the same timepoints than the swelling studies, namely day 0, 1, 3, 7, 10, 14, 21, and 28 after photopolymerization. Actual hydrogel thickness and diameter were recorded prior to compression. A 2.5 N load cell was used at a crosshead speed of 0.5 mm/min and samples were kept hydrated in DPBS at RT for the duration of the test. The Young’s modulus was calculated as the slope of the stress-strain curve in the linear range of 10-15% strain using **Equation 6**:

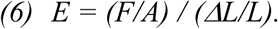

Three hydrogels per timepoint were measured and the experiments were repeated three times (n=3, N=9). The data were processed with a custom Python 3 script.

#### Stress-relaxation

Stress-relaxation studies were performed together with uniaxial compression tests. Samples were compressed until 15% strain as described in section above. The strain was then kept constant for 1000 seconds, while the stress was recorded.

#### Collagenase Assay

PVA-GEL hydrogels were immersed in 3 ml of DPBS and incubated at 37°C for 24 hours to allow them to reach equilibrium and wash off unreacted polymers. All samples were then removed from the DPBS, immersed in 3 ml of DPBS containing 3 mM of CaCl_2_ and 0.02 mg/ml of collagenase Type I (ThermoFisher Scientific, Cat. #17018029) and incubated at 37°C at different timepoints. Enzymatic degradation studies were performed at 0, 3, 6, 9, 12, 24, and 168 hours, after the hydrogels reached equilibrium. Samples were then removed from the collagenase solution, lyophilised, and weighted to determine their final dry mass. The residual weight was obtained by calculating the mass loss, as described in previous sections. Three samples were prepared for each timepoint and the experiments were repeated three times (n=3, N=9).

### Primary Cell Isolation and 3D Encapsulation

#### Primary Astrocyte Isolation

Primary astrocytes were obtained from the hippocampus of post-natal day 4 Sprague Dawley rat pups P4^106,107^. Pups were sacrificed with a lethal overdose of pentobarbital and decapitated with surgical scissors. The brain was removed from the skull and placed under a dissection microscope. The two hemispheres were separated, and the meninges and blood vessels were removed thoroughly to remove any contaminant cells. The hippocampus of each hemisphere was identified based on its characteriztic moon shaped morphology and separated from the cortex. The collected tissues were mechanically dissociated with a 1 ml pipette tip followed by a 23G needle. Single cells were counted and plated in poly-L-lysine (PLL)-coated T75 flasks at a density of 40000 cells/cm^2^ in plating medium (Dulbecco’s modified Eagle’s medium/F12 - Cat. # D6421 – supplemented with 33 mM D-glucose, 1% L-glutamine, 1% penicillin/streptomycin, 1% FBS, 2% B27). Flasks were incubated in a humidified atmosphere with 5% CO_2_ at 37°C, and the medium was changed every 2-3 days. After 10 days of culture, cells were shaken for 6 hours at 37°C at 200 rpm^108^. Flasks were washed twice to remove detached cells and then passaged. Enriched cultures of mature astrocytes were obtained after 2-3 weeks, and cells were used from P3 to P6.

#### Primary Astrocytes Encapsulation

Primary astrocytes were transitioned gradually from plating medium to serum free medium (Dulbecco’s modified Eagle’s medium/F12 - Cat. #D6421 – supplemented with 33 mM D-glucose, 1% L-glutamine, 1% penicillin/streptomycin, 1% BSA, 1% G5) the week before encapsulation. To encapsulate primary astrocytes in PVA-GEL hydrogels, cells were directly added to the macromer solution with a density of 5 million cells/ml (equal to 100,000 cells per hydrogel) and homogeneously mixed with a pipette. The cell suspension was poured into the PDMS disk moulds (6 mm diameter and 0.5 mm thickness, 20 μl each). Photopolymerization was carried out with visible light at 15 mW/cm^2^ for 3 min. Hydrogels were placed in 24 well plates and 750 µl of medium was added to each hydrogel. Hydrogels were then moved to an incubator (5% CO_2_ at 37°C). The day after, 500 µl of medium were removed and replaced with fresh serum free medium. Medium was refreshed every 2-3 days by replacing 500 µl of the medium with fresh serum free medium. Samples were kept in culture for 7 days (medium change schematic in Figure S5).

#### Primary VM cell Isolation

Primary mixed cultures were obtained from the VM of Sprague–Dawley rat embryos^77^. Tissue dissociation was performed as described by Thompson and Parish^109^. Briefly, the midbrain of E14 embrys was carefully dissected from the embryo and the neural tube was open to expose the ventral region. The dissected tissues were then incubated for 5 min at 37°C in 0.1% trypsin and mechanically dissociated with a 1 ml pipette tip, followed by a 23G needle. Single cells were then counted and plated in ultra-low adhesion T75 flasks at a density of 1.5 -2 x 10^5^ cells/ml in VM plating medium (Dulbecco’s modified Eagle’s medium/F12 - Cat. # D6421 - supplemented with 33 mM D-glucose, 1% L-glutamine, 1% penicillin/streptomycin, 1% FBS, 2% B27, 20 ng/ml EGF, 20 ng/ml FGF).

#### Primary Neurosphere Culture

Primary VM cells were cultured in ultra-low adhesion flasks to promote the growth of the isolated cell suspension as neurospheres. Neurosphere cultures were maintained *in vitro* for 7-10 days until an average diameter of 50-100 µm was achieved. The cultures were then gradually transitioned from VM plating medium to encapsulation medium (Dulbecco’s modified Eagle’s medium/F12 - Cat. # D6421 – supplemented with 33 mM D-glucose, 1% L-glutamine, 1% penicillin/streptomycin, 0.5% FBS, 1% B27, 10 ng/ml EGF, 10 ng/ml FGF, 50 µg/ml BSA, 0.5% N2) following a step-by-step approach shown in Figure S5 and the whole medium was replaced every 3-4 days.

#### Primary Astrocytes and Neurospheres Encapsulation

Co-cultures of primary mature astrocytes and VM neurospheres were encapsulated in PVA-GEL hydrogels by directly mixing the astrocytes and the neurospheres into the macromer solution (10 wt% nominal polymer content, 25:75 PVA:GEL). Primary astrocytes were encapsulated at a density of 5 x 10^6^ cells/ml (i.e., 1 x 105 cells per hydrogel), while VM neurospheres suspension was first passed through a 100-µm cell strainer and then added to the polymer solution at a density of 1 x 10^5^ neurospheres/ml (i.e., 2 x 10^3^ neurospheres per hydrogel). The hydrogel precursor was then poured into PDMS disk moulds (6 mm diameter and 0.5 mm thickness) and photocrosslinked with visible light at 15 mW/cm^2^ for 3 min, as described before. On the day after encapsulation, 500 µl of medium were replaced with fresh encapsulation medium. Afterwards, medium was refreshed every 2-3 days by replacing 500 µl of medium in order to gradually transition the cultures to serum-free medium (Figure S5). Samples were kept in culture for 7 or 14 days.

#### Brain Slice Dissection

Hippocampal slices were prepared on porous culture inserts following the interface method described by De Simoni and Yu^110^. Briefly, brains were bisected longitudinally to separate the hemispheres and then sectioned into 300-µm coronal slices using a 7000smz-2 Vibrotome (Campden Instruments). The hippocampal formation was identified and dissected from each slice under a stereomicroscope using straight ultra-fine forceps and placed into 12-mm PTFE Millicell culture inserts with a 0.4-µm pore size (Millipore, Cat. # PICM01250). Tlice culture medium was composed of MEM GlutaMAX-I (Gibco, Cat. # 41090036) supplemented with 25% horse serum (HS, Gibco, Cat. # 26050070), 23% EBSS, 36.11 mM D-glucose (Merck, Cat. # G5767), and 1% penicillin/streptomycin (Gibco, Cat. # 15070063). Half of the medium was refreshed every 3 days.

### Cell Assays and Functional Characterisation

#### Metabolic Activity

The metabolic activity of encapsulated primary astrocytes was evaluated using an Alamar Blue assay (ThermoFisher, DAL1025). Samples were incubated for 5 hours in the Alamar Blue solution (10% Alamar Blue assay in serum free medium). The supernatant was collected and analysed using a plate reader (544-590nm). The absorbance (A) values difference was calculated as a proxy for the total metabolic reduction in the well. The values were then normalized by the ratio between the difference in absorbance between the positive control (Alamar Blue-only well, AB) and negative controls (no Alamar Blue, C) at 540 nm and at 590 nm (**Equation 7**):

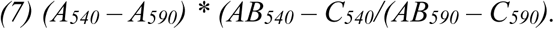

#### Gel Zymography

Gelatin zymography was performed to evaluate the production of the matrix metalloproteinase-2 (MMP-2) by encapsulated cells^111^. Briefly, the supernatant of encapsulated primary astrocytes was collected at DIV3, DIV7, and DIV10 post-encapsulation. The zymography resolving gel solution was prepared by mixing 4.6 ml DI water, 2.7 ml 30% acrylamide (Sigma A3699), 2.5 ml 1.5 M Tris (pH 8.8), 100 μl 10% sodium dodecyl sulphate (SDS, Sigma 1.06022), 285 μl 2.8 mg/ml gelatin (Sigma G2500) in DI water, 6 μl tetramethylethylenediamine (TEMED, Sigma T9281) and 100 μl 10% ammonium persulfate (APS, Sigma A3678). The zymography loading gel was prepared by adding 3.4 ml DI water, 830 μl 30% acrylamide, 630 μl 1 M Tris (pH 6.8), 50 μl 10% SDS, 5 μl TEMED and 50 μl 10% APS. The MMP-2 (Sigma PF037) standard was prepared at a concentration of 5 ng/ml and non-reducing Laemmli buffer (Thermo Fisher Scientific 84788) was added at a ratio of 1:3. Zymography samples were prepared by adding non-reducing Laemmli buffer to the collected supernatants (1:3). Zymography was run for 50 min at 200 V in Tris-Glycine SDS Running Buffer (10X) (Thermo Fisher Scientific LC2675). The gels were washed 4 times for 15 min with 2.5% (v/v) Triton X-100 with DI water. The gels were incubated for 15 hours at 37°C in the developing buffer (10 ml 1 M Tris (pH 7.5), 8 ml 5 M NaCl, 1 ml 1 M CaCl2, 1.6 ml 2.5% Triton X-100 and 179.4 ml DI water). Once developed, the gels were stained for 1 hour in Coomassie blue staining solution (0.5 g Brilliant Blue - Sigma 27816, 250 ml methanol, 100 ml acetic acid - Sigma A6283, and 150 ml DI water) and then destained for approximately 4 hours with destaining solution (150 ml methanol, 5 ml formic acid – Sigma F0507, and 350 ml DI water). Gels were imaged using a UVP Biospectrum 500 Imaging System and the bands were analysed using the ImageJ densitometry plugin. The amount of MMP-2 in the 3D samples was normalised by the total number of cells.

#### Immunofluorescent Staining

To assess the development of astrocytes in PVA-NB hydrogels, samples were stained via indirect double-immunofluorescent labelling of relevant biomarkers. Samples were first fixed with 4% PFA for 20 min at RT. Hydrogels were then washed twice in DPBS and incubated in permeabilization solution (0.5 ml Triton X-100, 10.3 g sucrose, 0.292 g NaCl, 0.06 g MgCl2, 0.476 g HEPES buffer, in 100 ml water, pH 7.2) for 5 min at RT. Non-specific binding sites were blocked with 1% BSA for 30 min at 37°C, and incubated at RT for 24 hours with a chicken anti-GFAP antibody (1:250, Abcam 4674), rabbit anti-paxillin antibody (1:200, Abcam 32084), and mouse anti-MMP-2 antibody (1:200, Santa Cruz sc-13594). Following incubation with the primary antibodies, samples were washed 3 times with 0.05% Tween 20/DPBS and incubated at RT overnight with Alexa Fluor 555 goat anti-Rabbit IgG (H+L) (1:100, ThermoFisher Scientific A-21429), Alexa Fluor 594 goat anti-Mouse IgG (H+L) (1:250, ThermoFisher Scientific A-11005), and Alexa Fluor 647 goat anti-Chicken IgY (H+L) (1:250, ThermoFisher Scientific A-21449) antibodies. After washing 3 times with DPBS, cell nuclei were counterstained with Hoechst 33342 (1:2000, ThermoFisher Scientific 62249).

For morphological and phenotypic assessment, the following primary antibodies were used: chicken anti-GFAP (1:1000, ab4674), rabbit anti-myelin PLP (1:200, ab254363), and mouse anti-βIII tubulin (1:500, ab78078). To evaluate the establishment of synaptic connections, the following primary antibodies were used: chicken anti-βIII tubulin (1:1000, ab41489), rabbit anti-synaptophysin (1:400, ab52636), and mouse anti-PSD95 (1:200, ab13552).

Organotypic cultures were processed for IFS following the protocol described by Gogolla et al^112^. After fixation, samples were washed once with cold DPBS, and then incubated in cold 20% MeOH DPBS for another 5 min. Samples were then washed and permeabilised by incubating in 0.5% Triton X-100 in DPBS overnight at 4°C. On the following day, samples were blocked with 20% BSA in DPBS for 4 hours at RT. Samples were then incubated overnight at 4°C with the primary antibodies solubilised in 5% BSA in DPBS. The following primary antibodies were used: chicken anti-GFAP (1:1000, ab4674), rabbit anti-paxillin (1:200, Abcam 32084), and mouse anti-βIII tubulin (1:500, ab78078). To evaluate the establishment of synaptic connections, the following primary antibodies were used: chicken anti-βIII tubulin (1:1000, ab41489), rabbit anti-synaptophysin (1:400, ab52636), and mouse anti-PSD95 (1:200, ab13552).

#### Calcium Imaging

Calcium Imaging was performed using a Fluo-4 AM assay (ThermoFisher Scientific F14201). Samples were washed once in DPBS and then incubated for 1 hour in the staining solution (1 µM Fluo-4 and 2.5 mM probenecid diluted in recording medium comprised of HBSS supplemented with 128 mM NaCl, 1 mM MgCl2, 45 mM sucrose, 10 mM glucose, and 0.01 M HEPES). Samples were washed once with fresh recording medium. Samples were then imaged at 37°C and 5% CO^2^ on a Zeiss Axio Observer widefield fluorescence microscope using a 10x air objective. Each area was imaged for 5 min with an acquisition rate of 8.5 Hz and at least three areas per sample were imaged.

#### Microscopy and Image Analysis

Hydrogels were imaged with a Leica SP8 inverted confocal microscope with a fixed scan of 1024×1024 pixels. On average, 3 z-stacked images at 63x magnification were acquired from each sample. The image analysis was performed using the ImageJ software (National Institutes of Health, United States). For primary astrocytes cultures, cell nuclei were used to count the number of cells per selected area. Cell elongation was determined from the GFAP expression (green channel) and cells were marked as “not elongated” if the assumed a round morphology, or as “elongated” if they showed extended processes. An example of a cell considered elongated is the 25:75 condition in **Figure 4D**. GFAP expression was calculated applying a threshold to all images, and the GFAP positive area was divided by the total number of cells to obtain the average GFAP expression per cell. Paxillin and MMP-2 expression were calculated applying a threshold to the images, and the resulting values were normalised to the GFAP expression. For co-cultures in PVA-GEL hydrogels and organotypic cultures, 3 z-stacked images at 20x or 63x magnification were acquired from each sample. Images were filtered to reduce the noise level using the functions ‘Remove Outliers’ and ‘Minimum’ filter from ImageJ. Relative biomarker expression (GFAP, βIII tubulin, myelin, synaptophysin, PSD-95) was determined by applying a threshold to all images and calculating the percentage of the entire area covered by a given fluorescence. The percentages of astrocytes and neurons were calculated based on the expression of GFAP and βIII tubulin normalised by the total GFAP and βIII tubulin expression per image (Figure S3).

Calcium imaging analysis was performed using the Intensity 2 function of the ImageJ software. Regions of interest (ROIs) were drawn on the image to select the cells to analyse, and the Intensity 2 function was used to detect changes in fluorescence intensity over time.

## Statistical analysis

All reported data represent the mean of 3 repeats (n=3), each of them consisting of at least 3 replicates (N=9). All results are expressed as the mean ± the standard deviation of the means. One-way ANOVA was performed to compare the means of the groups and a Tukey-Kramer test was used for multiple comparisons between the means. Differences were considered significant at a significance level of 5% (* p < 0.05), 1% (** p < 0.01) or 0.1% (*** p < 0.001).

## Acknowledgements

The authors acknowledge funding support from the European Research Council through the Consolidator Grant 771985. The authors thank Aaron Lee for initial support in the drafting of the manuscript. Axel Moore for the technical support with the compression experiments, Dariusz Lachowski for the experimental support with the Zymography experiments. We acknowledge the support of the technicians from the Bioengineering Department, and the Facility for Imaging by Light Microscopy (FILM) at the faculty of Medicine, Imperial College London.The authors declare no conflict of interest.

## 6. Data Availability Statement

Supplementary material can be found in the Supplementary Document S1 (containing Figure S1-S5 and Table S1-S8). All the original scripts used in this study has been deposited at Zenodo, which is publicly available as of the date of submission (doi: https://zenodo.org/records/16414597).

## References

1. Organic Neuroelectronics: From Neural Interfaces to Neuroprosthetics - Go - 2022 - Advanced Materials - Wiley Online Library. https://advanced.onlinelibrary.wiley.com/doi/10.1002/adma.202201864.

2. Sha, B. & Du, Z. Neural repair and regeneration interfaces: a comprehensive review. Biomed. Mater. 19, 022002 (2024).

3. Portillo-Lara, R., Goding, J. A. & Green, R. A. Adaptive biomimicry: design of neural interfaces with enhanced biointegration. Curr. Opin. Biotechnol. 72, 62–68 (2021).

4. Boulingre, M., Portillo-Lara, R. & A. Green, R. Biohybrid neural interfaces: improving the biological integration of neural implants. (2023) doi:10.1039/D3CC05006H.

5. Lotti, F., Ranieri, F., Vadalà, G., Zollo, L. & Di Pino, G. Invasive Intraneural Interfaces: Foreign Body Reaction Issues. Front. Neurosci. 11, (2017).

6. Ferguson, M., Sharma, D., Ross, D. & Zhao, F. A Critical Review of Microelectrode Arrays and Strategies for Improving Neural Interfaces. Adv. Healthc. Mater. 8, 1900558 (2019).

7. Gori, M., Vadalà, G., Giannitelli, S. M., Denaro, V. & Di Pino, G. Biomedical and Tissue Engineering Strategies to Control Foreign Body Reaction to Invasive Neural Electrodes. Front. Bioeng. Biotechnol. 9, (2021).

8. Szostak, K. M., Grand, L. & Constandinou, T. G. Neural Interfaces for Intracortical Recording: Requirements, Fabrication Methods, and Characteristics. Front. Neurosci. 11, (2017).

9. Boufidis, D., Garg, R., Angelopoulos, E., Cullen, D. K. & Vitale, F. Bio-inspired electronics: Soft, biohybrid, and “living” neural interfaces. Nat. Commun. 16, 1861 (2025).

10. Boulingre, M., Genta, M., Goding, J., Lara, R. P. & Green, R. Tissue-Engineered Interfaces to Enhance the Biointegration of Neural Implants. in *TISSUE ENGINEERING PART A* vol. 29 (MARY ANN LIEBERT, INC 140 HUGUENOT STREET, 3RD FL, NEW ROCHELLE, NY 10801 USA, 2023).

11. Goding, J. et al. A living electrode construct for incorporation of cells into bionic devices. MRS Commun. 7, 487–495 (2017).

12. Goding, J. A., Gilmour, A. D., Aregueta-Robles, U. A., Hasan, E. A. & Green, R. A. Living Bioelectronics: Strategies for Developing an Effective Long-Term Implant with Functional Neural Connections. Adv. Funct. Mater. 28, 1–20 (2018).

13. Rochford, A. E. et al. When Bio Meets Technology: Biohybrid Neural Interfaces. Adv. Mater. 32, 1903182 (2020).

14. George, J., Hsu, C. C., Nguyen, L. T. B., Ye, H. & Cui, Z. Neural tissue engineering with structured hydrogels in CNS models and therapies. Biotechnol. Adv. 42, 1–17 (2020).

15. Qazi, T. H. et al. Programming hydrogels to probe spatiotemporal cell biology. Cell Stem Cell 29, 678–691 (2022).

16. Madhusudanan, P., Raju, G. & Shankarappa, S. Hydrogel systems and their role in neural tissue engineering. J. R. Soc. Interface 17, 20190505 (2020).

17. Neural tissue engineering of the CNS using hydrogels - Nisbet - 2008 - Journal of Biomedical Materials Research Part B: Applied Biomaterials - Wiley Online Library. https://onlinelibrary.wiley.com/doi/full/10.1002/jbm.b.31000.

18. Salatino, J. W., Ludwig, K. A., Kozai, T. D. Y. & Purcell, E. K. Glial responses to implanted electrodes in the brain. Nat. Biomed. Eng. 2017 111 1, 862–877 (2017).

19. Adewole, D. O., Serruya, M. D., Wolf, J. A. & Cullen, D. K. Bioactive Neuroelectronic Interfaces. Front. Neurosci. 13, 1–11 (2019).

20. Chaudhuri, O., Cooper-White, J., Janmey, P. A., Mooney, D. J. & Shenoy, V. B. Effects of extracellular matrix viscoelasticity on cellular behaviour. Nature 584, 535–546 (2020).

21. Lam, D. et al. Tissue-specific extracellular matrix accelerates the formation of neural networks and communities in a neuron-glia co-culture on a multi-electrode array. Sci. Rep. 9, 4159 (2019).

22. Jovanov Milošević, N., Judaš, M., Aronica, E. & Kostovic, I. Neural ECM in laminar organization and connectivity development in healthy and diseased human brain. Prog. Brain Res. 214, 159–178 (2014).

23. Lu, P., Takai, K., Weaver, V. M. & Werb, Z. Extracellular Matrix Degradation and Remodeling in Development and Disease. Cold Spring Harb. Perspect. Biol. 3, a005058 (2011).

24. Miyata, S. & Kitagawa, H. Formation and remodeling of the brain extracellular matrix in neural plasticity: Roles of chondroitin sulfate and hyaluronan. Biochim. Biophys. Acta BBA - Gen. Subj. 1861, 2420–2434 (2017).

25. A glial perspective on the extracellular matrix and perineuronal net remodeling in the central nervous system. https://www.frontiersin.org/journals/cellular-neuroscience/articles/10.3389/fncel.2022.1022754/full.

26. Clarke, L. E. & Barres, B. A. Emerging roles of astrocytes in neural circuit development. Nat. Rev. Neurosci. 14, 311–321 (2013).

27. Dai, Y. et al. Effects and Mechanism of Action of Neonatal Versus Adult Astrocytes on Neural Stem Cell Proliferation After Traumatic Brain Injury. Stem Cells Dayt. Ohio 37, 1344– 1356 (2019).

28. Markey, K. M., Saunders, J. C., Smuts, J., von Reyn, C. R. & Garcia, A. D. R. Astrocyte development—More questions than answers. Front. Cell Dev. Biol. 11, (2023).

29. Wang, F. et al. Roles of activated astrocyte in neural stem cell proliferation and differentiation. Stem Cell Res. 7, 41–53 (2011).

30. Koshy, S. T. et al. Click-Crosslinked Injectable Gelatin Hydrogels. Adv. Healthc. Mater. 5, 541–547 (2016).

31. Hu, Y. et al. Matrix stiffness changes affect astrocyte phenotype in an in vitro injury model. NPG Asia Mater. 13, 35 (2021).

32. Galarza, S., Crosby, A. J., Pak, C. & Peyton, S. R. Control of Astrocyte Quiescence and Activation in a Synthetic Brain Hydrogel. Adv. Healthc. Mater. 9, 1901419 (2020).

33. Peters, S. B. Co-culture methods to study neuronal function and disease. Neural Regen. Res. 16, 972–973 (2020).

34. De Simone, U., Caloni, F., Gribaldo, L. & Coccini, T. Human Co-culture Model of Neurons and Astrocytes to Test Acute Cytotoxicity of Neurotoxic Compounds. Int. J. Toxicol. 36, 463–477 (2017).

35. Ioannou, M. S., Liu, Z. & Lippincott-Schwartz, J. A Neuron-Glia Co-culture System for Studying Intercellular Lipid Transport. Curr. Protoc. Cell Biol. 84, e95 (2019).

36. Luchena, C. et al. A Neuron, Microglia, and Astrocyte Triple Co-culture Model to Study Alzheimer’s Disease. Front. Aging Neurosci. 14, 844534 (2022).

37. Goshi, N., Morgan, R. K., Lein, P. J. & Seker, E. A primary neural cell culture model to study neuron, astrocyte, and microglia interactions in neuroinflammation. J. Neuroinflammation 17, 155 (2020).

38. Raimondi, I., Tunesi, M., Forloni, G., Albani, D. & Giordano, C. 3D brain tissue physiological model with co-cultured primary neurons and glial cells in hydrogels. J. Tissue Eng. 11, (2020).

39. Liu, R. et al. From 2D to 3D Co-Culture Systems: A Review of Co-Culture Models to Study the Neural Cells Interaction. Int. J. Mol. Sci. 23, 13116 (2022).

40. Dalrymple, A. N. et al. Electrochemical and biological performance of chronically stimulated conductive hydrogel electrodes. J. Neural Eng. 17, 026018 (2020).

41. Vallejo-Giraldo, C., Genta, M., Cauvi, O., Goding, J. & Green, R. Hydrogels for 3D Neural Tissue Models: Understanding Cell-Material Interactions at a Molecular Level. Front. Bioeng. Biotechnol. 8, (2020).

42. Van Hoorick, J. et al. Highly Reactive Thiol-Norbornene Photo-Click Hydrogels: Toward Improved Processability. Macromol. Rapid Commun. 39, 1–7 (2018).

43. Boulingre, M. et al. Multi-layered electrode constructs for neural tissue engineering. J. Mater. Chem. B 13, 3390–3404 (2025).

44. Chiareli, R. A. et al. The Role of Astrocytes in the Neurorepair Process. Front. Cell Dev. Biol. 9, 1–23 (2021).

45. Farhy-Tselnicker, I. & Allen, N. J. Astrocytes, neurons, synapses: a tripartite view on cortical circuit development. Neural Develop. 13, (2018).

46. Aregueta-Robles, U. A., Martens, P. J., Poole-Warren, L. A. & Green, R. A. Tailoring 3D hydrogel systems for neuronal encapsulation in living electrodes. J. Polym. Sci. Part B Polym. Phys. 56, 273–287 (2018).

47. Mũnoz, Z., Shih, H. & Lin, C. C. Gelatin hydrogels formed by orthogonal thiol-norbornene photochemistry for cell encapsulation. Biomater. Sci. 2, 1063–1072 (2014).

48. Lin, C.-C., Frahm, E. & Afolabi, F. O. Orthogonally Crosslinked Gelatin-Norbornene Hydrogels for Biomedical Applications. Macromol. Biosci. 24, 2300371 (2024).

49. Göckler, T. et al. Tuning Superfast Curing Thiol-Norbornene-Functionalized Gelatin Hydrogels for 3D Bioprinting. Adv. Healthc. Mater. 10, 2100206 (2021).

50. Qin, X. H., Wang, X., Rottmar, M., Nelson, B. J. & Maniura-Weber, K. Near-Infrared Light-Sensitive Polyvinyl Alcohol Hydrogel Photoresist for Spatiotemporal Control of Cell-Instructive 3D Microenvironments. Adv. Mater. 30, (2018).

51. Lim, K. S. et al. New Visible-Light Photoinitiating System for Improved Print Fidelity in Gelatin-Based Bioinks. ACS Biomater. Sci. Eng. 2, 1752–1762 (2016).

52. Shih, H. & Lin, C. C. Visible-Light-Mediated Thiol-Ene Hydrogelation Using Eosin-Y as the Only Photoinitiator. Macromol. Rapid Commun. 34, 269–273 (2013).

53. Summonte, S., Racaniello, G. F., Lopedota, A., Denora, N. & Bernkop-Schnürch, A. Thiolated polymeric hydrogels for biomedical application: Cross-linking mechanisms. J. Controlled Release 330, 470–482 (2021).

54. Lu, P. et al. Harnessing the potential of hydrogels for advanced therapeutic applications: current achievements and future directions. Signal Transduct. Target. Ther. 9, 166 (2024).

55. Chiarugi, P. & Giannoni, E. Anoikis: A necessary death program for anchorage-dependent cells. Biochem. Pharmacol. 76, 1352–1364 (2008).

56. Ishikawa, F., Ushida, K., Mori, K. & Shibanuma, M. Loss of anchorage primarily induces non-apoptotic cell death in a human mammary epithelial cell line under atypical focal adhesion kinase signaling. Cell Death Dis. 2015 61 6, e1619–e1619 (2015).

57. Tang, S., Richardson, B. M. & Anseth, K. S. Dynamic covalent hydrogels as biomaterials to mimic the viscoelasticity of soft tissues. Prog. Mater. Sci. 120, 100738 (2021).

58. Lou, J., Stowers, R., Nam, S., Xia, Y. & Chaudhuri, O. Stress relaxing hyaluronic acid-collagen hydrogels promote cell spreading, fiber remodeling, and focal adhesion formation in 3D cell culture. Biomaterials 154, 213–222 (2018).

59. Castrillón Martínez, D. C., Zuluaga, C. L., Restrepo-Osorio, A. & Álvarez-López, C. Characterization of sericin obtained from cocoons and silk yarns. Procedia Eng. 200, 377–383 (2017).

60. Kim, S., Nimni, M. E., Yang, Z. & Han, B. Chitosan/gelatin-based films crosslinked by proanthocyanidin. J. Biomed. Mater. Res. - Part B Appl. Biomater. 75, 442–450 (2005).

61. Thomas, M. & Willerth, S. M. 3-D bioprinting of neural tissue for applications in cell therapy and drug screening. Front. Bioeng. Biotechnol. 5, 69 (2017).

62. Anwar, M. et al. Formulation and evaluation of interpenetrating network of xanthan gum and polyvinylpyrrolidone as a hydrophilic matrix for controlled drug delivery system. Polym. Bull. 78, 59–80 (2021).

63. Parlato, M., Reichert, S., Barney, N. & Murphy, W. L. Poly(ethylene glycol) hydrogels with adaptable mechanical and degradation properties for use in biomedical applications. Macromol. Biosci. 14, 687–698 (2014).

64. Martínez-Cerdeño, V. & Noctor, S. C. Neural progenitor cell terminology. Front. Neuroanat. 12, 1–8 (2018).

65. Gilmour, A. D., Woolley, A. J., Poole-Warren, L. A., Thomson, C. E. & Green, R. A. A critical review of cell culture strategies for modelling intracortical brain implant material reactions. Biomaterials 91, 23–43 (2016).

66. Zhu, M. et al. Gelatin methacryloyl and its hydrogels with an exceptional degree of controllability and batch-to-batch consistency. Sci. Rep. 9, 1–13 (2019).

67. Sofia Peressotti, Gillian E. Koehl, Josef A. Goding, Rylie A. Green. Self-Assembling Hydrogel Structures for Neural Tissue Repair. ACS Biomater. Sci. Eng. 7, (2021).

68. Heath, D. E. & Cooper, S. L. The development of polymeric biomaterials inspired by the extracellular matrix. J. Biomater. Sci. Polym. Ed. 28, 1051–1069 (2017).

69. Guvendiren, M. & Burdick, J. A. Stiffening hydrogels to probe short- and long-term cellular responses to dynamic mechanics. Nat. Commun. 2012 31 3, 1–9 (2012).

70. Sagdic, K., Fernández-Lavado, E., Mariello, M., Akouissi, O. & Lacour, S. P. Hydrogels and conductive hydrogels for implantable bioelectronics. MRS Bull. 48, 495–505 (2023).

71. Chaudhuri, O. et al. Hydrogels with tunable stress relaxation regulate stem cell fate and activity. Nat. Mater. 15, 326–334 (2016).

72. Wei, Z., Schnellmann, R., Pruitt, H. C. & Gerecht, S. Hydrogel Network Dynamics Regulate Vascular Morphogenesis. Cell Stem Cell 27, 798–812.e6 (2020).

73. Nguyen, H. D., Sun, X., Yokota, H. & Lin, C. C. Probing Osteocyte Functions in Gelatin Hydrogels with Tunable Viscoelasticity. Biomacromolecules 22, 1115–1126 (2021).

74. Bertsch, P., Andrée, L., Besheli, N. H. & Leeuwenburgh, S. C. G. Colloidal hydrogels made of gelatin nanoparticles exhibit fast stress relaxation at strains relevant for cell activity. Acta Biomater. 138, 124–132 (2022).

75. Green, M. A., Bilston, L. E. & Sinkus, R. In vivo brain viscoelastic properties measured by magnetic resonance elastography. NMR Biomed. 21, 755–764 (2008).

76. Fan, C. et al. Effect of type-2 astrocytes on the viability of dorsal root ganglion neurons and length of neuronal processes. Neural Regen. Res. 9, 119 (2014).

77. Vallejo-Giraldo, C. et al. Attenuated Glial Reactivity on Topographically Functionalized Poly(3,4-Ethylenedioxythiophene):P-Toluene Sulfonate (PEDOT:PTS) Neuroelectrodes Fabricated by Microimprint Lithography. Small 14, 1800863 (2018).

78. Fang, A. et al. Effects of astrocyte on neuronal outgrowth in a layered 3D structure. Biomed. Eng. Online 18, 1–16 (2019).

79. Hopkins, A. M., DeSimone, E., Chwalek, K. & Kaplan, D. L. 3D in vitro modeling of the central nervous system. Prog. Neurobiol. 125, 1–25 (2015).

80. Becerra-Calixto, A. & Cardona-Gómez, G. P. The role of astrocytes in neuroprotection after brain stroke: Potential in cell therapy. Front. Mol. Neurosci. 10, 88 (2017).

81. Huang, Z. et al. YAP stabilizes SMAD1 and promotes BMP2-induced neocortical astrocytic differentiation. Development 143, 2398–2409 (2016).

82. Oschmann, F., Berry, H., Obermayer, K. & Lenk, K. From in silico astrocyte cell models to neuron-astrocyte network models: A review. Brain Res. Bull. 136, 76–84 (2018).

83. Karahuseyinoglu, S. et al. Three-dimensional neuron–astrocyte construction on matrigel enhances establishment of functional voltage-gated sodium channels. J. Neurochem. 156, 848– 866 (2021).

84. DePalma, T. J. et al. Tuning a bioengineered hydrogel for studying astrocyte reactivity in glioblastoma. Acta Biomater. 189, 155–167 (2024).

85. Kim, B. J. et al. Astrocyte-Encapsulated Hydrogel Microfibers Enhance Neuronal Circuit Generation. Adv. Healthc. Mater. 9, e1901072 (2020).

86. Matthiesen, I. et al. Astrocyte 3D culture and bioprinting using peptide functionalized hyaluronan hydrogels. Sci. Technol. Adv. Mater. 24, 2165871 (2023).

87. Chan, S. J. et al. Promoting Neuro-Supportive Properties of Astrocytes with Epidermal Growth Factor Hydrogels. Stem Cells Transl. Med. 8, 1242–1248 (2019).

88. Yao, L., Sai, H. V., Shippy, T. & Li, B. Cellular and Transcriptional Response of Human Astrocytes to Hybrid Protein Materials. ACS Appl. Bio Mater. 7, 2887–2898 (2024).

89. Gao, Y.-Y. et al. Gelatin-Based Hydrogel for Three-Dimensional Neuron Culture Application. ACS Omega 8, 45288–45300 (2023).

90. Lim, K. S., Alves, M. H., Poole-Warren, L. A. & Martens, P. J. Covalent incorporation of non-chemically modified gelatin into degradable PVA-tyramine hydrogels. Biomaterials 34, 7097–7105 (2013).

91. Kang, A. R., Park, J. S., Ju, J., Jeong, G. S. & Lee, S. H. Cell encapsulation via microtechnologies. Biomaterials 35, 2651–2663 (2014).

92. Ingavle, G. C., Gehrke, S. H. & Detamore, M. S. The bioactivity of agarose-PEGDA interpenetrating network hydrogels with covalently immobilized RGD peptides and physically entrapped aggrecan. Biomaterials 35, 3558–3570 (2014).

93. Klotz, B. J. et al. A Versatile Biosynthetic Hydrogel Platform for Engineering of Tissue Analogues. Adv. Healthc. Mater. 8, (2019).

94. Agrawal, S. M., Lau, L. & Yong, V. W. MMPs in the central nervous system: Where the good guys go bad. Semin. Cell Dev. Biol. 19, 42–51 (2008).

95. Ethell, I. M. & Ethell, D. W. Matrix metalloproteinases in brain development and remodeling: Synaptic functions and targets. J. Neurosci. Res. 85, 2813–2823 (2007).

96. Sarem, M. et al. Interplay between stiffness and degradation of architectured gelatin hydrogels leads to differential modulation of chondrogenesis in vitro and in vivo. Acta Biomater. 69, 83–94 (2018).

97. Stogsdill, J. A., Harwell, C. C. & Goldman, S. A. Astrocytes as master modulators of neural networks: Synaptic functions and disease-associated dysfunction of astrocytes. Ann. N. Y. Acad. Sci. 1525, 41–60 (2023).

98. Herculano-Houzel, S. The glia/neuron ratio: How it varies uniformly across brain structures and species and what that means for brain physiology and evolution. Glia 62, 1377–1391 (2014).

99. Van Horn, M. R. & Ruthazer, E. S. Glial regulation of synapse maturation and stabilization in the developing nervous system. Curr. Opin. Neurobiol. 54, 113–119 (2019).

100. Tibbitt, M. W., Kloxin, A. M., Sawicki, L. A. & Anseth, K. S. Mechanical Properties and Degradation of Chain and Step-Polymerized Photodegradable Hydrogels. Macromolecules 46, 2785–2792 (2013).

101. Zustiak, S. P., Pubill, S., Ribeiro, A. & Leach, J. B. Hydrolytically degradable poly(ethylene glycol) hydrogel scaffolds as a cell delivery vehicle: Characterization of PC12 cell response. Biotechnol. Prog. 29, 1255–1264 (2013).

102. Stoppini, L., Buchs, P.-A. & Muller, D. A simple method for organotypic cultures of nervous tissue. J. Neurosci. Methods 37, 173–182 (1991).

103. Sifringer, L. et al. An Implantable Biohybrid Neural Interface Toward Synaptic Deep Brain Stimulation. Adv. Funct. Mater. 35, 2416557 (2025).

104. Zhu, R. et al. Electrical stimulation affects neural stem cell fate and function in vitro. Exp. Neurol. 319, 112963 (2019).

105. Chen, C., Bai, X., Ding, Y. & Lee, I.-S. Electrical stimulation as a novel tool for regulating cell behavior in tissue engineering. Biomater. Res. 23, 25 (2019).

106. Czupalla, C. J., Yousef, H., Wyss-Coray, T. & Butcher, E. C. Collagenase-based Single Cell Isolation of Primary Murine Brain Endothelial Cells Using Flow Cytometry. Bio-Protoc. 8, (2018).

107. O’Sullivan, S. A., O’Sullivan, C., Healy, L. M., Dev, K. K. & Sheridan, G. K. Sphingosine 1-phosphate receptors regulate TLR4-induced CXCL5 release from astrocytes and microglia. J. Neurochem. 144, 736–747 (2018).

108. Schildge, S., Bohrer, C., Beck, K. & Schachtrup, C. Isolation and culture of mouse cortical astrocytes. J. Vis. Exp. JoVE (2013) doi:10.3791/50079.

109. Thompson, L. H. & Parish, C. L. Transplantation of fetal midbrain dopamine progenitors into a rodent model of Parkinson’s disease. Methods Mol. Biol. 1059, 169–180 (2013).

110. De Simoni, A. & MY Yu, L. Preparation of organotypic hippocampal slice cultures: interface method. Nat. Protoc. 1, 1439–1445 (2006).

111. Lachowski, D. et al. Matrix stiffness modulates the activity of MMP-9 and TIMP-1 in hepatic stellate cells to perpetuate fibrosis. Sci. Rep. 2019 91 9, 1–9 (2019).

112. Gogolla, N., Galimberti, I., DePaola, V. & Caroni, P. Preparation of organotypic hippocampal slice cultures for long-term live imaging. Nat. Protoc. 2006 13 1, 1165–1171 (2006).

