## Supplementary Document S1 for "Astrocyte-Guided Maturation of Neural Constructs in a Modular Biosynthetic Hydrogel for Biohybrid Neurotechnologies"

### Supplementary Information

**Title:** Harnessing astrocytes for neural support: a tailorable, bio-orthogonal hydrogel for functional brain interfaces.

*Martina Genta, Sofia Peressotti, Roberto Portillo-Lara, Josef Goding, Rylie Green**

**A**

| Peak | Peak Location (δ) | Integral | Norbornene groups per PVA macromer |
|---|---|---|---|
| A | 4 | 6.51 | 363.2 |
| E | 6.2 | 1.00 | 27.9 |

| Groups | Peak Location (δ) | Number of hydrogens | Integral | Mol per 100 g of gelatin |
|---|---|---|---|---|
| Phenol | 7.3 | 5 | 1.00 | 0.0155 [297] |
| Norbornene | 6.0-6.3 | 2 | 0.62 | 0.024 |

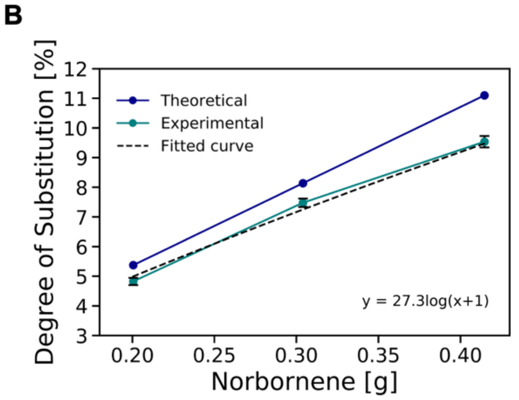


**Figure S1. Degrees of substitution of the PVA-NB and GEL-NB, related to Figure 2.** (A) Above: Integral values of characteristic peaks from PVA-NB $^1$H NMR spectroscopy. Integral values were normalized to the E peak. Below: integral values of characteristic peaks from GEL-NB $^1$H NMR spectroscopy. Integral values were normalized to the phenol peak. (B) Comparison between the theorical amount of norbornene needed to reach a given DS and the experimental values obtained. The DS of GEL-NB was calculated by comparing the number of moles of NB crosslinked to the GEL backbone to the number of moles of carboxyl groups (COOH) present in the GEL molecule (0.1267 mol/100 g of gelatin), obtaining an actual DS of 18.9% (i.e., 0.024 / 0.1267).

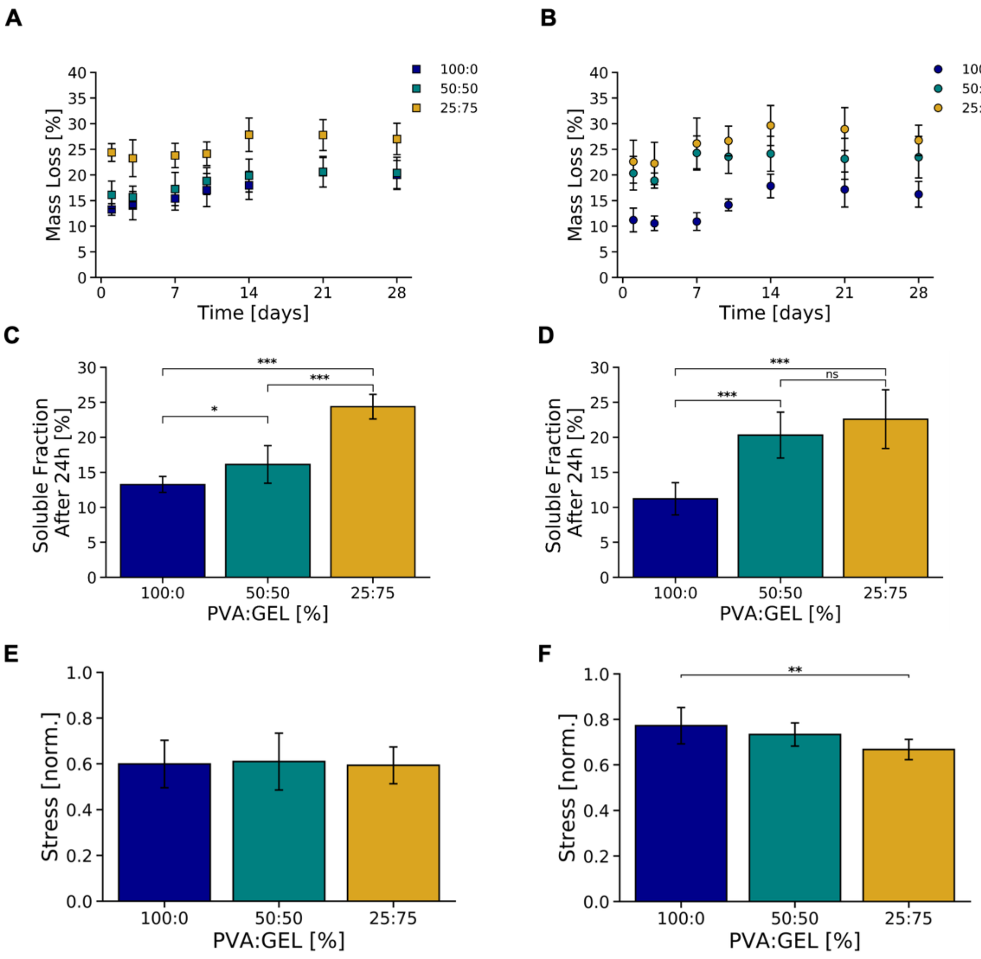


**Figure S2. Additional physical and mechanical characterizations of the PVA-GEL hydrogel, related to Figure 3 and Figure 4.** (A-B) Overall mass loss of (A) 10 wt% PVA-GEL hydrogels and (B) 7.5 wt% PVA-GEL hydrogels with different polymer ratios. (C-D) Soluble fraction of (C) 10 wt% and (D) 7.5 wt% PVA-GEL hydrogels with different polymer ratios after 24 hours of incubation in DPBS at 37◦C. (E-F) Stress-relaxation behaviour of PVA-GEL hydrogels after 24 hours of incubation in DPBS. Relaxation of (E) 10 wt% and (F) 7.5 wt% PVA-GEL hydrogels reached after 15 min of constant strain. All reported data represent the mean of 3 repeats (n=3), each of them consisting of at least 3 replicates (N=9). All results are expressed as the mean ± the standard deviation of the means. One-way ANOVA was performed to compare the means of the groups and a Tukey-Kramer test was used for multiple comparisons between the means. Differences were considered significant at a significance level of 5% (* $p < 0.05$), 1% (** $p < 0.01$) or 0.1% (*** $p < 0.001$). Statistical analyses results shown in **Table S3, S4**.

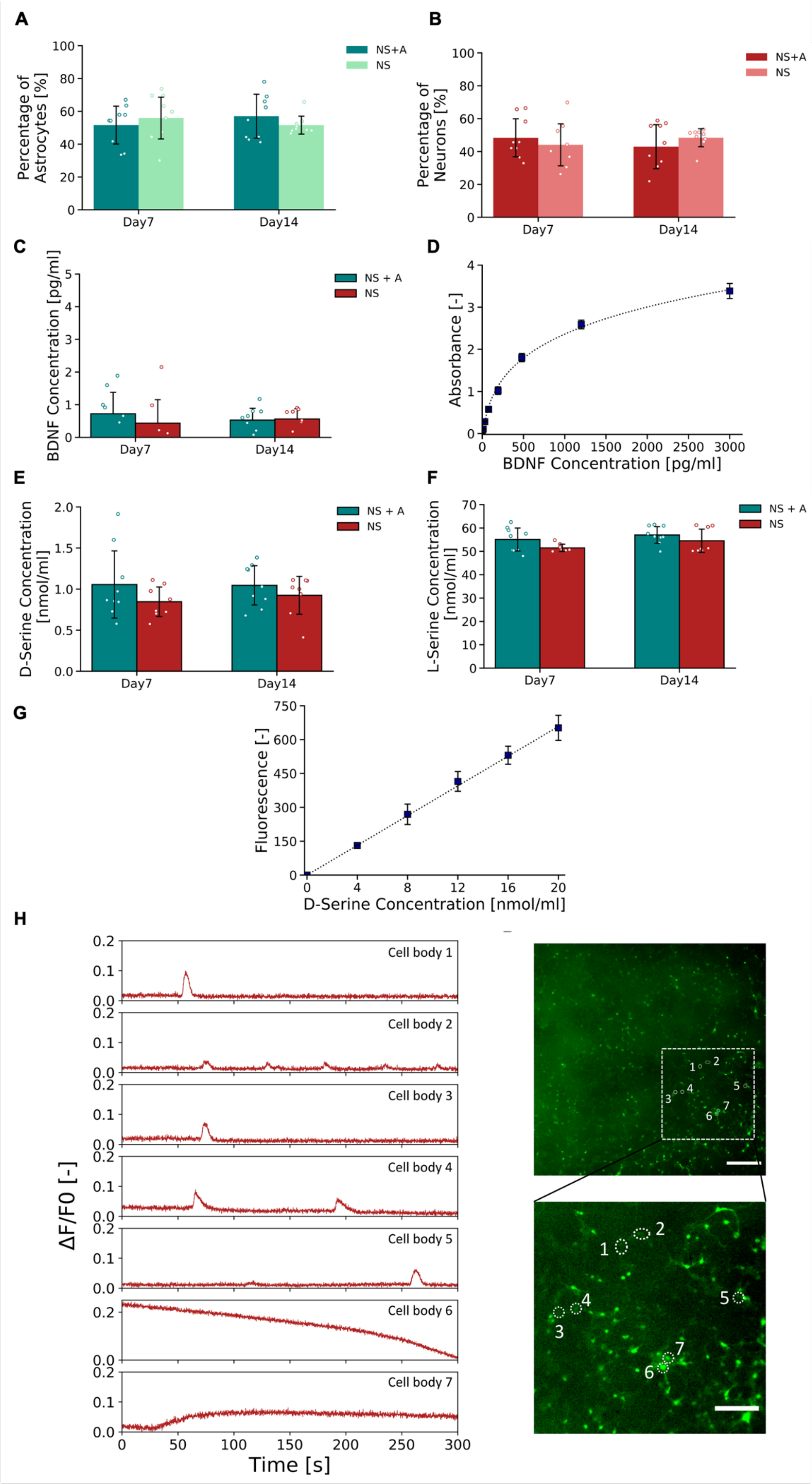

A
B
C
D
E
F
G
H
Percentage of Astrocytes [%]
Percentage of Neurons [%]
NS+A
NS
Day7
Day14
BDNF Concentration [pg/ml]
NS + A
Absorbance [-]
BDNF Concentration [pg/ml]
D-Serine Concentration [nmol/ml]
L-Serine Concentration [nmol/ml]
Fluorescence [-]
D-Serine Concentration [nmol/ml]
ΔF/F0 [-]
Cell body 1
Cell body 2
Cell body 3
Cell body 4
Cell body 5
Cell body 6
Cell body 7
Time [s]

**Figure S3. Neuro-glia culture evaluation, related to Figure 7.** (A-B) Percentage of (A) astrocytes and (B) neurons encapsulated in PVA-GEL hydrogels at DIV7 and DIV14. n = 3. (C) Quantification of BDNF secreted by cell-laden PVA-GEL hydrogels at DIV7 and DIV14. (D) Standard curve for the BDNF ELISA assay. (E-F) Quantitative analyses of the concentration of (E) D-Serine and (F) L-Serine secreted by different cell cultures in PVA-GEL hydrogels at DIV7 and DIV14. n = 3. (G) Standard curve with increasing D-Serine concentrations. (H) Functional characterization of neural network activity in cell-laden PVA-GEL hydrogels at DIV7. Calcium activity was recorded from co-cultures of NS and astrocytes encapsulated in PVA-GEL hydrogels at DIV14. Representative ROIs containing the recorded cells surrounded by white dotted lines are shown in the immunofluorescence panel. Scale bar = 200 μm.

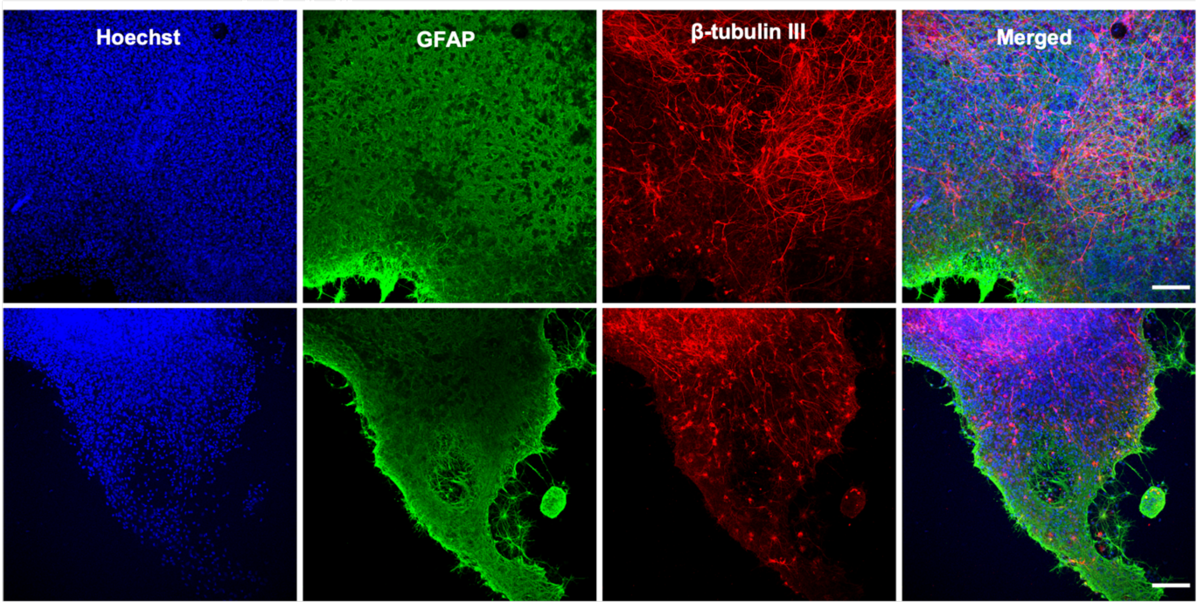


**Figure S4. Representative fluorescent micrographs of organotypic hippocampal slice cultures at DIV14, related to Figure 8.** (A) Cell nuclei are shown in blue, (B) GFAP+ astrocytes are shown in green and (C) β-tubulin III+ neurons in red. Images with all fluorescent channels merged are shown in (D). Scale bar = 100 μm.

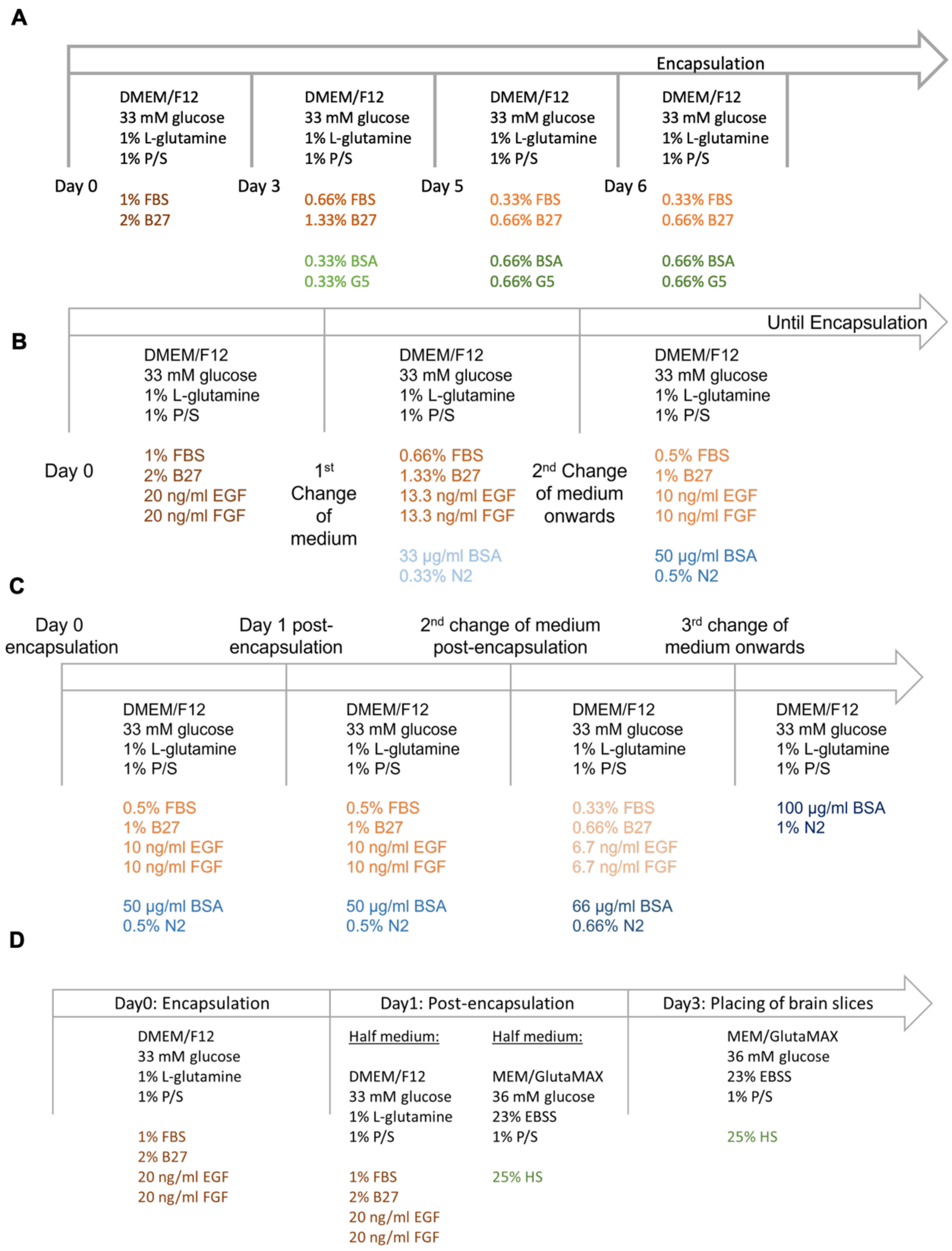


**Figure S5. Schematics of medium formulations, related to Experimental Section.** (A) In vitro culture scheme for primary astrocytes. Transitioned from plating medium to serum free medium. (B) In vitro culture scheme for primary NS showing the transition from plating medium to encapsulation medium. (C) In vitro culture scheme for primary astrocyte and NS co-cultures encapsulated in PVA-GEL hydrogels showing the transition from encapsulation medium to serum-free medium. (D) In vitro culture scheme for organotypic cultures showing the transition from VM plating medium to slice culture medium.

**Table S1:** Statistical testing of 10 wt% PVA-GEL hydrogels - Swelling studies - *p <0.05, **p <0.01, **p <0.001, n=3. Related to **Figure 3**.

| Condition 1 (PVA:GEL) | Condition 2 (PVA:GEL) | P Value | Significance |
|---|---|---|---|
| 100:0 d1 | 100:0 d3 | 0.900 | - |
| 100:0 d3 | 100:0 d7 | 0.900 | - |
| 100:0 d7 | 100:0 d10 | 0.650 | - |
| 100:0 d10 | 100:0 d14 | 0.770 | - |
| 100:0 d14 | 100:0 d21 | 0.001 | *** |
| 100:0 d21 | 100:0 d28 | 0.001 | *** |
| 100:0 d1 | 100:0 d28 | 0.001 | *** |
| 50:50 d1 | 50:50 d3 | 0.900 | - |
| 50:50 d3 | 50:50 d7 | 0.900 | - |
| 50:50 d7 | 50:50 d10 | 0.900 | - |
| 50:50 d10 | 50:50 d14 | 0.900 | - |
| 50:50 d14 | 50:50 d21 | 0.900 | - |
| 50:50 d21 | 50:50 d28 | 0.900 | - |
| 50:50 d1 | 50:50 d28 | 0.900 | - |
| 25:75 d1 | 25:75 d3 | 0.900 | - |
| 25:75 d3 | 25:75 d7 | 0.900 | - |
| 25:75 d7 | 25:75 d10 | 0.900 | - |
| 25:75 d10 | 25:75 d14 | 0.900 | - |
| 25:75 d14 | 25:75 d21 | 0.900 | - |
| 25:75 d21 | 25:75 d28 | 0.900 | - |
| 25:75 d1 | 25:75 d28 | 0.900 | - |
| 100:0 d1 | 25:75 d1 | 0.001 | *** |
| 100:0 d1 | 50:50 d1 | 0.001 | *** |
| 25:75 d1 | 50:50 d1 | 0.900 | - |
| 100:0 d3 | 25:75 d3 | 0.001 | *** |
| 100:0 d3 | 50:50 d3 | 0.001 | *** |
| 25:75 d3 | 50:50 d3 | 0.900 | - |
| 100:0 d7 | 25:75 d7 | 0.001 | *** |
| 100:0 d7 | 50:50 d7 | 0.001 | *** |
| 25:75 d7 | 50:50 d7 | 0.900 | - |
| 100:0 d10 | 25:75 d10 | 0.001 | *** |
| 100:0 d10 | 50:50 d10 | 0.001 | *** |
| 25:75 d10 | 50:50 d10 | 0.900 | - |
| 100:0 d14 | 25:75 d14 | 0.001 | *** |
| 100:0 d14 | 50:50 d14 | 0.001 | *** |
| 25:75 d14 | 50:50 d14 | 0.900 | - |
| 100:0 d21 | 25:75 d21 | 0.001 | *** |
| 100:0 d21 | 50:50 d21 | 0.001 | *** |
| 25:75 d21 | 50:50 d21 | 0.900 | - |
| 100:0 d28 | 25:75 d28 | 0.001 | *** |
| 100:0 d28 | 50:50 d28 | 0.001 | *** |
| 25:75 d28 | 50:50 d28 | 0.704 | - |

**Table S2**: Statistical testing of 7.5 wt% PVA-GEL hydrogels - Swelling studies - *p <0.05, **p <0.01, **p <0.001, n=3. Related to **Figure 3**.

| Condition 1 (PVA:GEL) | Condition 2 (PVA:GEL) | P Value | Significance |
|---|---|---|---|
| 100:0 d1 | 100:0 d3 | 0.900 | - |
| 100:0 d3 | 100:0 d7 | 0.900 | - |
| 100:0 d7 | 100:0 d10 | 0.900 | - |
| 100:0 d10 | 100:0 d14 | 0.315 | - |
| 100:0 d14 | 100:0 d21 | 0.169 | - |
| 100:0 d21 | 100:0 d28 | 0.001 | *** |
| 100:0 d1 | 100:0 d28 | 0.001 | *** |
| 50:50 d1 | 50:50 d3 | 0.900 | - |
| 50:50 d3 | 50:50 d7 | 0.900 | - |
| 50:50 d7 | 50:50 d10 | 0.900 | - |
| 50:50 d10 | 50:50 d14 | 0.900 | - |
| 50:50 d14 | 50:50 d21 | 0.900 | - |
| 50:50 d21 | 50:50 d28 | 0.900 | - |
| 50:50 d1 | 50:50 d28 | 0.900 | - |
| 25:75 d1 | 25:75 d3 | 0.900 | - |
| 25:75 d3 | 25:75 d7 | 0.900 | - |
| 25:75 d7 | 25:75 d10 | 0.900 | - |
| 25:75 d10 | 25:75 d14 | 0.900 | - |
| 25:75 d14 | 25:75 d21 | 0.900 | - |
| 25:75 d21 | 25:75 d28 | 0.900 | - |
| 25:75 d1 | 25:75 d28 | 0.900 | - |
| 100:0 d1 | 25:75 d1 | 0.001 | *** |
| 100:0 d1 | 50:50 d1 | 0.001 | *** |
| 25:75 d1 | 50:50 d1 | 0.900 | - |
| 100:0 d3 | 25:75 d3 | 0.001 | *** |
| 100:0 d3 | 50:50 d3 | 0.001 | *** |
| 25:75 d3 | 50:50 d3 | 0.900 | - |
| 100:0 d7 | 25:75 d7 | 0.001 | *** |
| 100:0 d7 | 50:50 d7 | 0.001 | *** |
| 25:75 d7 | 50:50 d7 | 0.900 | - |
| 100:0 d10 | 25:75 d10 | 0.001 | *** |
| 100:0 d10 | 50:50 d10 | 0.001 | *** |
| 25:75 d10 | 50:50 d10 | 0.900 | - |
| 100:0 d14 | 25:75 d14 | 0.001 | *** |
| 100:0 d14 | 50:50 d14 | 0.001 | *** |
| 25:75 d14 | 50:50 d14 | 0.900 | - |
| 100:0 d21 | 25:75 d21 | 0.001 | *** |
| 100:0 d21 | 50:50 d21 | 0.001 | *** |
| 25:75 d21 | 50:50 d21 | 0.900 | - |
| 100:0 d28 | 25:75 d28 | 0.001 | *** |
| 100:0 d28 | 50:50 d28 | 0.001 | *** |
| 25:75 d28 | 50:50 d28 | 0.496 | - |

**Table S3**: Statistical testing of 10 wt% PVA-GEL hydrogels - Mass loss studies - *p <0.05, **p <0.01, **p <0.001, n=3. Related to **Figure S1**.

| Condition 1 (PVA:GEL) | Condition 2 (PVA:GEL) | P Value | Significance |
|---|---|---|---|
| 100:0 d1 | 100:0 d3 | 0.900 | - |
| 100:0 d3 | 100:0 d7 | 0.900 | - |
| 100:0 d7 | 100:0 d10 | 0.900 | - |
| 100:0 d10 | 100:0 d14 | 0.900 | - |
| 100:0 d14 | 100:0 d21 | 0.900 | - |
| 100:0 d21 | 100:0 d28 | 0.900 | - |
| 100:0 d1 | 100:0 d28 | 0.001 | *** |
| 50:50 d1 | 50:50 d3 | 0.900 | - |
| 50:50 d3 | 50:50 d7 | 0.900 | - |
| 50:50 d7 | 50:50 d10 | 0.900 | - |
| 50:50 d10 | 50:50 d14 | 0.900 | - |
| 50:50 d14 | 50:50 d21 | 0.900 | - |
| 50:50 d21 | 50:50 d28 | 0.900 | - |
| 50:50 d1 | 50:50 d28 | 0.141 | - |
| 25:75 d1 | 25:75 d3 | 0.900 | - |
| 25:75 d3 | 25:75 d7 | 0.900 | - |
| 25:75 d7 | 25:75 d10 | 0.900 | - |
| 25:75 d10 | 25:75 d14 | 0.365 | - |
| 25:75 d14 | 25:75 d21 | 0.900 | - |
| 25:75 d21 | 25:75 d28 | 0.900 | - |
| 25:75 d1 | 25:75 d28 | 0.900 | - |
| 100:0 d1 | 25:75 d1 | 0.001 | *** |
| 100:0 d1 | 50:50 d1 | 0.796 | - |
| 25:75 d1 | 50:50 d1 | 0.001 | *** |
| 100:0 d3 | 25:75 d3 | 0.001 | *** |
| 100:0 d3 | 50:50 d3 | 0.900 | - |
| 25:75 d3 | 50:50 d3 | 0.001 | *** |
| 100:0 d7 | 25:75 d7 | 0.001 | *** |
| 100:0 d7 | 50:50 d7 | 0.900 | - |
| 25:75 d7 | 50:50 d7 | 0.001 | *** |
| 100:0 d10 | 25:75 d10 | 0.001 | *** |
| 100:0 d10 | 50:50 d10 | 0.900 | - |
| 25:75 d10 | 50:50 d10 | 0.011 | * |
| 100:0 d14 | 25:75 d14 | 0.001 | *** |
| 100:0 d14 | 50:50 d14 | 0.900 | - |
| 25:75 d14 | 50:50 d14 | 0.001 | *** |
| 100:0 d21 | 25:75 d21 | 0.001 | *** |
| 100:0 d21 | 50:50 d21 | 0.900 | - |
| 25:75 d21 | 50:50 d21 | 0.001 | *** |
| 100:0 d28 | 25:75 d28 | 0.001 | *** |
| 100:0 d28 | 50:50 d28 | 0.900 | - |
| 25:75 d28 | 50:50 d28 | 0.001 | *** |

**Table S4**: Statistical testing of 7.5 wt% PVA-GEL hydrogels - Mass loss studies - *p <0.05, **p <0.01, **p <0.001, n=3. Related to **Figure S1**.

| Condition 1 (PVA:GEL) | Condition 2 (PVA:GEL) | P Value | Significance |
|---|---|---|---|
| 100:0 d1 | 100:0 d3 | 0.900 | - |
| 100:0 d3 | 100:0 d7 | 0.900 | - |
| 100:0 d7 | 100:0 d10 | 0.838 | - |
| 100:0 d10 | 100:0 d14 | 0.654 | - |
| 100:0 d14 | 100:0 d21 | 0.900 | - |
| 100:0 d21 | 100:0 d28 | 0.900 | - |
| 100:0 d1 | 100:0 d28 | 0.136 | - |
| 50:50 d1 | 50:50 d3 | 0.900 | - |
| 50:50 d3 | 50:50 d7 | 0.070 | - |
| 50:50 d7 | 50:50 d10 | 0.900 | - |
| 50:50 d10 | 50:50 d14 | 0.900 | - |
| 50:50 d14 | 50:50 d21 | 0.900 | - |
| 50:50 d21 | 50:50 d28 | 0.900 | - |
| 50:50 d1 | 50:50 d28 | 0.887 | - |
| 25:75 d1 | 25:75 d3 | 0.900 | - |
| 25:75 d3 | 25:75 d7 | 0.564 | - |
| 25:75 d7 | 25:75 d10 | 0.900 | - |
| 25:75 d10 | 25:75 d14 | 0.900 | - |
| 25:75 d14 | 25:75 d21 | 0.900 | - |
| 25:75 d21 | 25:75 d28 | 0.900 | - |
| 25:75 d1 | 25:75 d28 | 0.464 | - |
| 100:0 d1 | 25:75 d1 | 0.001 | *** |
| 100:0 d1 | 50:50 d1 | 0.001 | *** |
| 25:75 d1 | 50:50 d1 | 0.900 | - |
| 100:0 d3 | 25:75 d3 | 0.001 | *** |
| 100:0 d3 | 50:50 d3 | 0.001 | *** |
| 25:75 d3 | 50:50 d3 | 0.794 | - |
| 100:0 d7 | 25:75 d7 | 0.001 | *** |
| 100:0 d7 | 50:50 d7 | 0.001 | *** |
| 25:75 d7 | 50:50 d7 | 0.900 | - |
| 100:0 d10 | 25:75 d10 | 0.001 | *** |
| 100:0 d10 | 50:50 d10 | 0.001 | *** |
| 25:75 d10 | 50:50 d10 | 0.900 | - |
| 100:0 d14 | 25:75 d14 | 0.001 | *** |
| 100:0 d14 | 50:50 d14 | 0.010 | ** |
| 25:75 d14 | 50:50 d14 | 0.054 | - |
| 100:0 d21 | 25:75 d21 | 0.001 | *** |
| 100:0 d21 | 50:50 d21 | 0.022 | * |
| 25:75 d21 | 50:50 d21 | 0.030 | * |
| 100:0 d28 | 25:75 d28 | 0.001 | *** |
| 100:0 d28 | 50:50 d28 | 0.001 | *** |
| 25:75 d28 | 50:50 d28 | 0.827 | - |

**Table S5:** Statistical testing of 10 wt% PVA-GEL hydrogels - Mass loss studies normalised - *p <0.05, **p <0.01, **p <0.001, n=3. Related to **Figure 3**.

| Condition 1 (PVA:GEL) | Condition 2 (PVA:GEL) | P Value | Significance |
|---|---|---|---|
| 100:0 d1 | 100:0 d3 | 0.900 | - |
| 100:0 d3 | 100:0 d7 | 0.900 | - |
| 100:0 d7 | 100:0 d10 | 0.900 | - |
| 100:0 d10 | 100:0 d14 | 0.900 | - |
| 100:0 d14 | 100:0 d21 | 0.900 | - |
| 100:0 d21 | 100:0 d28 | 0.900 | - |
| 100:0 d1 | 100:0 d28 | 0.001 | *** |
| 50:50 d1 | 50:50 d3 | 0.900 | - |
| 50:50 d3 | 50:50 d7 | 0.900 | - |
| 50:50 d7 | 50:50 d10 | 0.900 | - |
| 50:50 d10 | 50:50 d14 | 0.900 | - |
| 50:50 d14 | 50:50 d21 | 0.900 | - |
| 50:50 d21 | 50:50 d28 | 0.900 | - |
| 50:50 d1 | 50:50 d28 | 0.141 | - |
| 25:75 d1 | 25:75 d3 | 0.900 | - |
| 25:75 d3 | 25:75 d7 | 0.900 | - |
| 25:75 d7 | 25:75 d10 | 0.900 | - |
| 25:75 d10 | 25:75 d14 | 0.365 | - |
| 25:75 d14 | 25:75 d21 | 0.900 | - |
| 25:75 d21 | 25:75 d28 | 0.900 | - |
| 25:75 d1 | 25:75 d28 | 0.900 | - |
| 100:0 d1 | 25:75 d1 | 0.900 | - |
| 100:0 d1 | 50:50 d1 | 0.900 | - |
| 25:75 d1 | 50:50 d1 | 0.900 | - |
| 100:0 d3 | 25:75 d3 | 0.900 | - |
| 100:0 d3 | 50:50 d3 | 0.900 | - |
| 25:75 d3 | 50:50 d3 | 0.900 | - |
| 100:0 d7 | 25:75 d7 | 0.866 | - |
| 100:0 d7 | 50:50 d7 | 0.900 | - |
| 25:75 d7 | 50:50 d7 | 0.900 | - |
| 100:0 d10 | 25:75 d10 | 0.217 | - |
| 100:0 d10 | 50:50 d10 | 0.900 | - |
| 25:75 d10 | 50:50 d10 | 0.765 | - |
| 100:0 d14 | 25:75 d14 | 0.900 | - |
| 100:0 d14 | 50:50 d14 | 0.900 | - |
| 25:75 d14 | 50:50 d14 | 0.900 | - |
| 100:0 d21 | 25:75 d21 | 0.288 | - |
| 100:0 d21 | 50:50 d21 | 0.837 | - |
| 25:75 d21 | 50:50 d21 | 0.900 | - |
| 100:0 d28 | 25:75 d28 | 0.194 | - |
| 100:0 d28 | 50:50 d28 | 0.900 | - |
| 25:75 d28 | 50:50 d28 | 0.900 | - |

**Table S6:** Statistical testing of 7.5 wt% PVA-GEL hydrogels - Mass loss studies normalised - *p <0.05, **p <0.01, **p <0.001, n=3. Related to **Figure 3**.

| Condition 1 (PVA:GEL) | Condition 2 (PVA:GEL) | P Value | Significance |
|---|---|---|---|

| | | | |
|---|---|---|---|
| 100:0 d1 | 100:0 d3 | 0.900 | - |
| 100:0 d3 | 100:0 d7 | 0.900 | - |
| 100:0 d7 | 100:0 d10 | 0.838 | - |
| 100:0 d10 | 100:0 d14 | 0.654 | - |
| 100:0 d14 | 100:0 d21 | 0.900 | - |
| 100:0 d21 | 100:0 d28 | 0.900 | - |
| 100:0 d1 | 100:0 d28 | 0.136 | - |
| 50:50 d1 | 50:50 d3 | 0.900 | - |
| 50:50 d3 | 50:50 d7 | 0.070 | - |
| 50:50 d7 | 50:50 d10 | 0.900 | - |
| 50:50 d10 | 50:50 d14 | 0.900 | - |
| 50:50 d14 | 50:50 d21 | 0.900 | - |
| 50:50 d21 | 50:50 d28 | 0.900 | - |
| 50:50 d1 | 50:50 d28 | 0.887 | - |
| 25:75 d1 | 25:75 d3 | 0.900 | - |
| 25:75 d3 | 25:75 d7 | 0.564 | - |
| 25:75 d7 | 25:75 d10 | 0.900 | - |
| 25:75 d10 | 25:75 d14 | 0.900 | - |
| 25:75 d14 | 25:75 d21 | 0.900 | - |
| 25:75 d21 | 25:75 d28 | 0.900 | - |
| 25:75 d1 | 25:75 d28 | 0.464 | - |
| 100:0 d1 | 25:75 d1 | 0.900 | - |
| 100:0 d1 | 50:50 d1 | 0.900 | - |
| 25:75 d1 | 50:50 d1 | 0.900 | - |
| 100:0 d3 | 25:75 d3 | 0.900 | - |
| 100:0 d3 | 50:50 d3 | 0.900 | - |
| 25:75 d3 | 50:50 d3 | 0.900 | - |
| 100:0 d7 | 25:75 d7 | 0.900 | - |
| 100:0 d7 | 50:50 d7 | 0.410 | - |
| 25:75 d7 | 50:50 d7 | 0.900 | - |
| 100:0 d10 | 25:75 d10 | 0.900 | - |
| 100:0 d10 | 50:50 d10 | 0.900 | - |
| 25:75 d10 | 50:50 d10 | 0.900 | - |
| 100:0 d14 | 25:75 d14 | 0.900 | - |
| 100:0 d14 | 50:50 d14 | 0.900 | - |
| 25:75 d14 | 50:50 d14 | 0.836 | - |
| 100:0 d21 | 25:75 d21 | 0.900 | - |
| 100:0 d21 | 50:50 d21 | 0.874 | - |
| 25:75 d21 | 50:50 d21 | 0.713 | - |
| 100:0 d28 | 25:75 d28 | 0.900 | - |
| 100:0 d28 | 50:50 d28 | 0.900 | - |
| 25:75 d28 | 50:50 d28 | 0.900 | - |

**Table S7:** Statistical testing of 10 wt% PVA-GEL hydrogels - Compression studies - *p <0.05, **p <0.01, **p <0.001, n=3. Related to **Figure 4**.

| Condition 1 (PVA:GEL) | Condition 2 (PVA:GEL) | P Value | Significance |
|---|---|---|---|
| 100:0 d0 | 100:0 d1 | 0.001 | *** |
| 100:0 d0 | 100:0 d28 | 0.001 | *** |
| 25:75 d0 | 25:75 d1 | 0.900 | - |
| 25:75 d0 | 25:75 d28 | 0.035 | * |

| | | | |
|---|---|---|---|
| 50:50 d0 | 50:50 d1 | 0.900 | - |
| 50:50 d0 | 50:50 d28 | 0.001 | *** |
| 100:0 d0 | 25:75 d0 | 0.001 | *** |
| 100:0 d0 | 50:50 d0 | 0.001 | *** |
| 25:75 d0 | 50:50 d0 | 0.001 | *** |
| 100:0 d1 | 25:75 d1 | 0.001 | *** |
| 100:0 d1 | 50:50 d1 | 0.900 | - |
| 25:75 d1 | 50:50 d1 | 0.001 | *** |
| 100:0 d3 | 25:75 d3 | 0.869 | - |
| 100:0 d3 | 50:50 d3 | 0.001 | *** |
| 25:75 d3 | 50:50 d3 | 0.001 | *** |
| 100:0 d7 | 25:75 d7 | 0.900 | - |
| 100:0 d7 | 50:50 d7 | 0.055 | - |
| 25:75 d7 | 50:50 d7 | 0.001 | *** |
| 100:0 d10 | 25:75 d10 | 0.900 | - |
| 100:0 d10 | 50:50 d10 | 0.208 | - |
| 25:75 d10 | 50:50 d10 | 0.026 | * |
| 100:0 d14 | 25:75 d14 | 0.900 | - |
| 100:0 d14 | 50:50 d14 | 0.083 | - |
| 25:75 d14 | 50:50 d14 | 0.175 | - |
| 100:0 d21 | 25:75 d21 | 0.900 | - |
| 100:0 d21 | 50:50 d21 | 0.001 | *** |
| 25:75 d21 | 50:50 d21 | 0.002 | ** |
| 100:0 d28 | 25:75 d28 | 0.886 | - |
| 100:0 d28 | 50:50 d28 | 0.001 | *** |
| 25:75 d28 | 50:50 d28 | 0.006 | ** |

**Table S8:** Statistical testing of 7.5 wt% PVA-GEL hydrogels - Compression studies - *p <0.05, **p <0.01, **p <0.001, n=3. Related to **Figure 4**.

| Condition 1 (PVA:GEL) | Condition 2 (PVA:GEL) | P Value | Significance |
|---|---|---|---|
| 100:0 d0 | 100:0 d1 | 0.075 | - |
| 100:0 d0 | 100:0 d28 | 0.001 | *** |
| 25:75 d0 | 25:75 d1 | 0.900 | - |
| 25:75 d0 | 25:75 d28 | 0.900 | - |
| 50:50 d0 | 50:50 d1 | 0.900 | - |
| 50:50 d0 | 50:50 d28 | 0.669 | - |
| 100:0 d0 | 25:75 d0 | 0.001 | *** |
| 100:0 d0 | 50:50 d0 | 0.001 | *** |
| 25:75 d0 | 50:50 d0 | 0.003 | ** |
| 100:0 d1 | 25:75 d1 | 0.001 | *** |
| 100:0 d1 | 50:50 d1 | 0.586 | - |
| 25:75 d1 | 50:50 d1 | 0.001 | *** |
| 100:0 d3 | 25:75 d3 | 0.001 | *** |
| 100:0 d3 | 50:50 d3 | 0.900 | - |
| 25:75 d3 | 50:50 d3 | 0.001 | *** |
| 100:0 d7 | 25:75 d7 | 0.001 | *** |
| 100:0 d7 | 50:50 d7 | 0.900 | - |
| 25:75 d7 | 50:50 d7 | 0.001 | *** |
| 100:0 d10 | 25:75 d10 | 0.003 | ** |
| 100:0 d10 | 50:50 d10 | 0.900 | - |

| | | | |
|---|---|---|---|
| 25:75 d10 | 50:50 d10 | 0.018 | * |
| 100:0 d14 | 25:75 d14 | 0.900 | - |
| 100:0 d14 | 50:50 d14 | 0.671 | - |
| 25:75 d14 | 50:50 d14 | 0.217 | - |
| 100:0 d21 | 25:75 d21 | 0.900 | - |
| 100:0 d21 | 50:50 d21 | 0.388 | - |
| 25:75 d21 | 50:50 d21 | 0.632 | - |
| 100:0 d28 | 25:75 d28 | 0.900 | - |
| 100:0 d28 | 50:50 d28 | 0.367 | - |
| 25:75 d28 | 50:50 d28 | 0.900 | - |
